# Nicotine engages a VTA-NAc feedback loop to inhibit amygdala-projecting dopamine neurons and induce anxiety

**DOI:** 10.1101/2024.08.20.607920

**Authors:** T. Le Borgne, C. Nguyen, E. Vicq, J. Jehl, C. Solié, N. Guyon, L. Daussy, A. Gulmez, L.M. Reynolds, S. Mondoloni, S. Tolu, S. Pons, U. Maskos, E. Valjent, A. Mourot, P. Faure, F. Marti

**Affiliations:** Plasticité du Cerveau CNRS UMR8249, École supérieure de physique et de chimie industrielles de la Ville de Paris (ESPCI Paris), Paris, France; Neuroscience Paris Seine CNRS UMR 8246 INSERM U1130, Institut de Biologie Paris Seine, Sorbonne Université, Paris, France; Université Paris Cité, CNRS, Unité de Biologie Fonctionnelle et Adaptative, F-75013, Paris, France; Institut Pasteur, Unité Neurobiologie intégrative des systèmes cholinergiques, Département de neuroscience, Paris, France; IGF, Université Montpellier, CNRS, Inserm, F-34094 Montpellier, France

**Keywords:** nicotine, dopamine circuits, ventral tegmental area, amygdala, nucleus accumbens, anxiety, nicotinic acetylcholine receptor, alcohol, addiction, in vivo electrophysiology

## Abstract

Nicotine activates ventral tegmental area (VTA) dopaminergic (DA) neurons projecting to the nucleus accumbens (NAc) to drive its reinforcing effects. Simultaneously, it inhibits those projecting to the amygdala (Amg) to mediate anxiety through a process that remains unknown. Here we show that NAc- and Amg-projecting DA neurons respond with similar polarities to ethanol and nicotine, suggesting a shared network-based mechanism underlying the inhibitory effect of these otherwise pharmacologically-distinct drugs. Selective activation of NAc-projecting DA neurons, using genetic or optogenetic strategies, produced inhibition of Amg-projecting DA neurons, through a GABAergic feedback loop. Furthermore, optogenetically silencing this feedback loop prevented nicotine from inducing both inhibition of DA neurons and anxiety-like behavior. Therefore, nicotine-induced inhibition of the VTA-Amg DA pathway results from a VTA-NAc inhibitory feedback loop, mediating anxiety.

## INTRODUCTION

Dopamine (DA) neurons in the ventral tegmental area (VTA) of the reward system play a crucial role in addiction mechanisms. Drugs of abuse, including nicotine and alcohol, act on this reward system through a variety of cellular mechanisms, ultimately leading to DA release in the nucleus accumbens (NAc) and behavioral reinforcement^1–3^. Indeed, global optogenetic activation of VTA DA neurons can induce conditioned place preference^4^ or reinforcement in a self-administration paradigm^5^, whereas inhibition of the same neurons induces avoidance behavior^6^. However, VTA DA neurons exhibit remarkable heterogeneity in their molecular properties and projection areas^7–9^. These neurons not only encode appetitive stimuli and reward predictions, but also salient signals such as aversive or alarming events^10–12^. Understanding the heterogeneity of drug effects on these neurons is therefore important to grasp the multifaceted nature of drug abuse.

Nicotine acts on nicotinic acetylcholine receptors (nAChRs), a family of ligand-gated ion channels^13^, and increases the activity of both DA and GABA neurons in the VTA^3,14^. Our previous work has shown that nicotine not only activates one population of VTA DA neurons as traditionally described, but also induces inhibition of another population of VTA DA neurons^15,16^. VTA DA neurons activated by nicotine project to the NAc and their optogenetic activation induces reinforcement, while VTA DA neurons inhibited by nicotine project to the amygdala (Amg) and their optogenetic inhibition mediates anxiety^16^. VTA DA neurons can thus mediate both the rewarding and anxiogenic properties of nicotine. Understanding how neurons projecting to Amg are inhibited by nicotine is, however, a major challenge in studying the dual effects of nicotine on reward and emotional circuits. We have previously shown a complete abolition of nicotine responses - both activation and inhibition - in mice deleted for the β2 nAChR subunit (β2^-/-^ mice), and restoration of both types of response following constitutive re-expression of the β2 subunit in the VTA (i.e. on all neuronal populations)^16^. These results indicate that nicotine activation and inhibition both stem from nicotine’s local action in the VTA. However, since nAChRs are cation-conducting ion channels, nicotine exerts a depolarising action on neurons expressing these receptors. It is therefore unlikely that nicotine directly inhibits any neuronal population. The observed nicotine-induced inhibition most likely involves a network effect arising from the activation of VTA cells expressing β2-containing nAChRs (β2*nAChRs).

Interestingly, alcohol, like nicotine, induces both activation or inhibition of distinct neuronal populations in the VTA^17^, suggesting that this pattern of opposite responses is not restricted to nicotine and may result from a circuit-based mechanism. Despite employing different molecular mechanisms^18^, these two drugs are known to increase DA release into the NAc by enhancing the activity of VTA DA neurons^16,19^. However, it remains unknown whether alcohol, like nicotine, inhibits a specific DA pathway, and whether these two drug-induced inhibitions share common features. In the present study, we investigated the mechanism by which nicotine and alcohol inhibit specific VTA DA neurons and induce anxiety.

## RESULTS

### Both nicotine and alcohol induce excitation and inhibition of distinct VTA DA neuron subpopulations *in vivo*

We first investigated whether nicotine and alcohol, two drugs with different molecular modes of action, could elicit similar responses in VTA DA neurons. To this end, *in vivo* single-cell juxtacellular recordings and labelling were performed in anesthetized mice to record the activity of VTA DA neurons during consecutive intravenous (i.v.) injection of nicotine (Nic; 30 µg/kg) and ethanol (EtOH; 250 mg/kg). All recorded neurons were confirmed as DA neurons by post hoc immunofluorescence with co-labeling for tyrosine hydroxylase and neurobiotin (TH+, NB+; **Figure 1A**). Acute i.v. paired injections of nicotine and ethanol induced a significant increase or decrease in the firing rate (**Figure 1B),** illustrated by bimodal distribution of firing frequency variations across neurons that was absent in vehicle injections (**Figure S1-C).** Among the 72 neurons recorded, 69 exhibited a significant response to nicotine while 67 neurons showed a significant response to ethanol compared to baseline variation (see Methods). Consistent with our previous work^16^, nicotine induced activation (Nic+; n = 41) or inhibition (Nic-; n = 28) of different populations of DA neurons (**Figure 1C)**. Similarly, neurons were activated (EtOH+, n = 52) or inhibited (EtOH-, n = 15) by ethanol (**Figure 1D**). Moreover, when comparing the individual responses of each neuron to nicotine and ethanol injections (**Figure S1D**), we found that 80% of the neurons responded with similar polarity to both substances (i.e., activated or inhibited by both drugs; **Figure S1E**). Strikingly, all neurons inhibited by ethanol were also inhibited by nicotine (15/15), suggesting that the two drugs act in a similar manner on these neurons. Furthermore, ethanol-activated and ethanol-inhibited neurons were anatomically segregated along the medio-lateral axis across the VTA (**Figure S1F**), as previously observed for nicotine^16^. Finally, the polarity of ethanol-induced responses was not dose-dependent, with neurons maintaining their activation or inhibition across all ethanol doses tested (**Figure S1G**). These results suggest that nicotine and ethanol induce inhibition in the same subpopulation of DA neurons.

**Figure 1:**
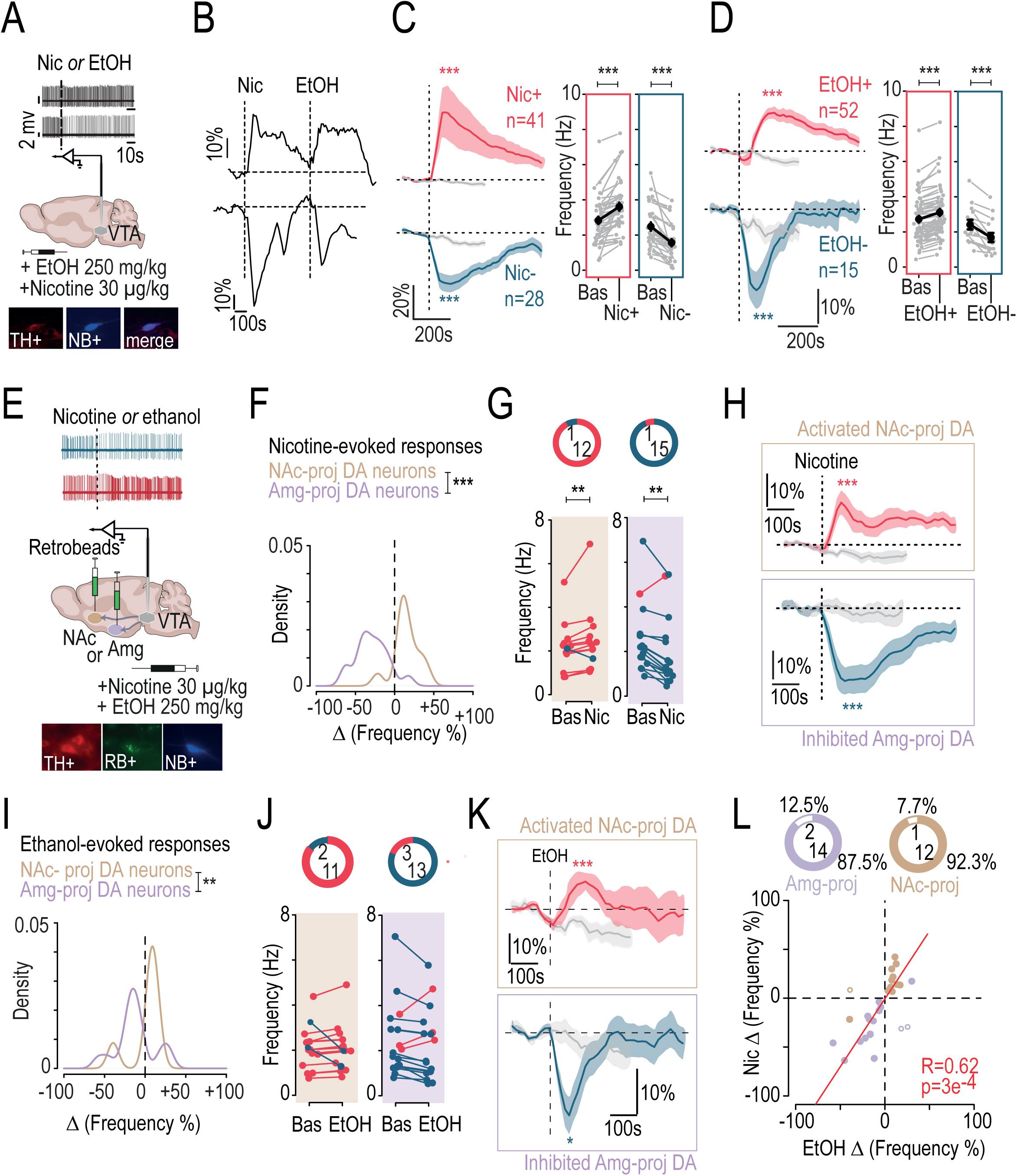
Nicotine and ethanol induce similar response profiles across VTA dopaminergic subpopulations. **(A)** In vivo juxtacellular recordings of VTA DA neuron responses to an intravenous (i.v.) injection of ethanol (EtOH; 250 mg/kg) or nicotine (Nic; 30 μg/kg). After recording, VTA DA neurons labeled with neurobiotin (NB) were subsequently identified by immunofluorescence of tyrosine hydroxylase (TH) and streptavidin-AMCA targeting neurobiotin (NB). **(B)** Examples of change in firing rate variation (% of baseline) after paired i.v. injection of nicotine and ethanol for two VTA DA neurons, one activated (*top*) and one inhibited (*bottom*) by the two drugs. **(C)** Left: time course of the mean change in firing frequency (% of baseline) after nicotine or saline (gray) i.v. injection in activated (Nic+; red; n = 41) and inhibited (Nic-; blue; n = 28) VTA DA neurons. Nicotine induced a significant increase or decrease in firing frequency compared to saline (paired Wilcoxon test: Nic+ vs Sal, V = 854, ***p < 0.001; Nic-vs Sal, V = 3, ***p < 0.001). Right: comparison of firing rate variation (Hz) between baseline and nicotine injection. Maximum firing rate after i.v. nicotine for Nic+ neurons or minimum firing rate after i.v. nicotine for Nic-neurons (paired Wilcoxon test, Nic+ vs Bas, V = 861, ***p < 0.001; Nic-vs Bas, V = 0, ***p < 0.001). **(D)** Same as (C) after ethanol or saline (gray) i.v. injection in activated (EtOH+, red, n = 52) and inhibited (EtOH-, blue, n = 15) VTA DA neurons (paired Wilcoxon test, EtOH+ vs Sal, V = 1290, ***p < 0.001; EtOH-vs Sal, V = 3, ***p < 0.001; EtOH+ vs Bas, V = 1378, ***p < 0.001; EtOH-vs Bas, V = 0, ***p < 0.001). **(E)** Retrobeads were injected either in the nucleus accumbens (NAc; injection in the lateral shell [NAcLSh] + medial shell [NAcMSh] + core [NAcCore]) or in the amygdala (Amg; injection in the basolateral [BLA] + central [CeA] nuclei) and *in vivo* recordings of VTA DA responses to an i.v. ethanol and nicotine injection were performed in anesthetized mice. Post hoc identification of NAc- or Amg-projecting DA neurons was performed by immunofluorescent co-labeling of tyrosine hydroxylase (TH), neurobiotin (NB), and retrobeads (RB). **(F)** Density of responses evoked by i.v. injections of nicotine in NAc- or Amg-projecting VTA DA neurons (brown, n = 13; purple, n = 16). Responses are expressed as a percentage of the change in firing frequency induced by the injection (Kolmogorov-Smirnov test, D = 0.86, ***p < 0.001). **(G)** Left: number of activated or inhibited VTA DA neurons (*top*) and firing rate variation between baseline and nicotine injection in NAc-projecting cells (*bottom*). Right: number of activated or inhibited VTA DA neurons (*top*) and firing rate variation between baseline and nicotine injection in Amg-projecting cells (*bottom*; paired Wilcoxon test: for NAc-proj, V = 80, **p = 0.01; for Amg-proj, V = 11, **p = 0.002). **(H)** Time course of the mean change in firing frequency (% of baseline) after i.v. injection of nicotine or saline (gray) for activated VTA DA neurons projecting to the NAc (*top*, n = 12) and inhibited VTA DA neurons projecting to the Amg (*bottom*, n = 15). Nicotine induced a significant increase or decrease in firing frequency compared to saline (paired Student t-test: for NAc-proj, t_11_ = 4.58, ***p < 0.001; for Amg-proj, t_13_ = -6.26, ***p < 0.001). **(I, J)** Same as (F, G) after i.v. injection of ethanol in NAc- or Amg-projecting VTA DA neurons (Kolmogorov-Smirnov test, D = 0.66, **p = 0.002). **(K)** Time course of mean change in firing frequency (% of baseline) after i.v. injection of ethanol or saline (gray) for activated VTA DA neurons projecting to the NAc (*top*, n = 11) and inhibited VTA DA neurons projecting to the Amg (*bottom*, n = 13). Ethanol induced a significant increase or decrease in firing frequency compared to saline (paired Student t-test: for NAc-proj, t_11_ = 4.83, ***p < 0.001; for Amg-proj, t_12_ = -2.34, *p = 0.04). **(L)** Correlation between ethanol- and nicotine-induced responses based on percentage of drug-induced frequency change for NAc- and Amg-projecting VTA DA neurons (Pearson’s correlation: t_27_ = 4.15, R = 0.62, ***p < 0.001). Percentage and number of correlated (firing rate decrease or increase for both drugs, filled dot) and uncorrelated (firing rate increase for one drug but decrease for the other, empty dot) response to nicotine or ethanol for NAc- and Amg-projecting VTA DA neurons (brown, n = 12/1; purple, n = 14/2).

### Nicotine and alcohol activate VTA DA neurons projecting to the NAc, while they inhibit those projecting to the Amg

To confirm that inhibited and activated neurons belong to distinct DA pathways, we injected retrobeads (RB) into either the Amg (BLA, CeA; **Figure S2A, B**) or the NAc (NAcLSh, NAcMSh, NAcCore; **Figure S2C, D**), followed by *in vivo* single-cell juxtacellular recordings during paired injection of the two drugs. Triple immunofluorescence labeling enabled post hoc confirmation of the DA nature (TH+) and the projection site (RB+; **Figure 1E**). As expected, anatomical segregation along the medio-lateral axis was observed between NAc- and Amg-projecting DA neurons (**Figure S2E**). Consistent with previous findings^16^, NAc-projecting DA neurons were mostly activated (12/13) whereas Amg-projecting DA neurons were predominantly inhibited (15/16), as indicated by the distribution of maximum frequency variation from baseline (**Figure 1F, G)** and the mean response relative to i.v. saline injection (**Figure 1H**). Intravenous ethanol injection produced a similar dichotomy in responses between NAc-projecting and Amg-projecting VTA DA neurons (**Figure 1I**). Most of the NAc-projecting DA neurons were activated (11/13), while most of the Amg-projecting DA neurons were inhibited (13/16) by i.v. ethanol injection (**Figure 1J, K**). Finally, the polarity of frequency variations induced by the two drugs was correlated in 87.5% of Amg-projecting and in 92.3% of NAc-projecting VTA DA neurons (**Figure 1L**). This indicates that nicotine and ethanol produce similar patterns of activation and inhibition in these distinct dopaminergic pathways. Specifically, it suggests a common network mechanism by which both drugs inhibit Amg-projecting dopamine neurons.

### NAc-projecting DA neurons are more sensitive to nicotine *ex vivo*

To understand the mechanism underlying the inhibition, we investigated the balance between direct excitatory and indirect inhibitory effects of nicotine on DA neurons projecting to the NAc (LSh) or the Amg (BLA) using patch-clamp recordings in VTA slices. We first examined nicotine-induced excitatory currents by applying nicotine puffs to neurons identified by co-labeling of TH, RB and biocytin (**Figure 2A**). Nicotine induced smaller currents in Amg-projecting DA neurons compared to NAc-projecting DA neurons at all doses tested (10, 30 and 100 µM; **Figure 2B**). This indicates that NAc-projecting DA neurons, which are known to be activated by nicotine *in vivo*, are more responsive to nicotine than Amg-projecting DA neurons, and suggests a differential expression of nAChRs in these two neuronal subpopulations.

**Figure 2:**
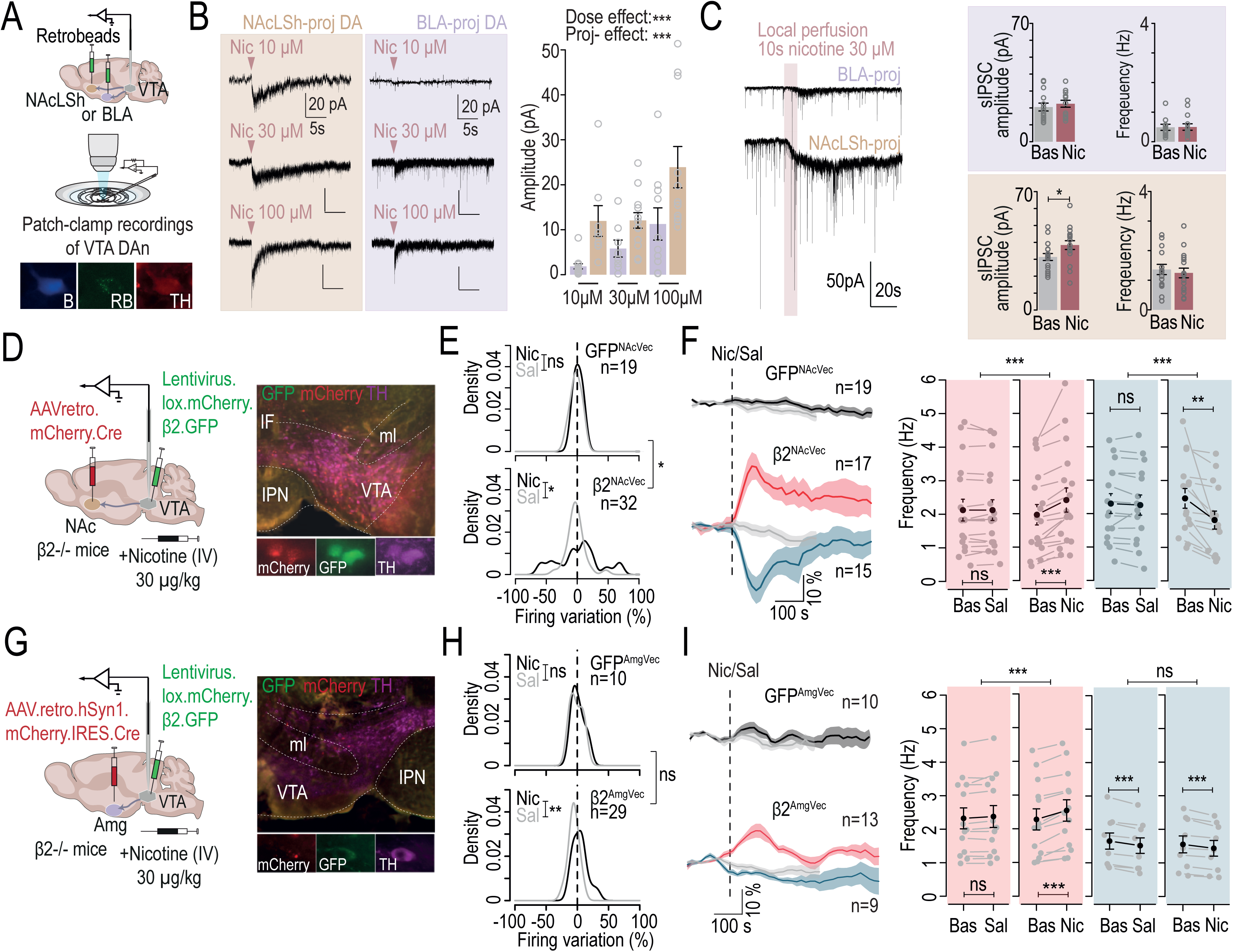
Nicotine-induced activation of NAc-projecting neurons triggers inhibition in the VTA. **(A)** Retrobeads were injected either in the NAcLSh or in the BLA and *ex vivo* patch-clamp recordings of VTA DA neurons were performed. Post hoc identification of NAc- or Amg-projecting DA neurons was performed by immunofluorescent co-labeling of tyrosine hydroxylase (TH, red), neurobiotin (NB, blue), and retrobeads (RB). **(B)** Left: example electrophysiological traces of nicotinic currents in NAcLSh-projecting or BLA-projecting DA neurons at three nicotine concentrations: 10, 30 and 100 μM (*left*). Right: mean currents evoked by local puffs of nicotine in either NAcLSh-projecting (brown; dose 10 µM, n = 8; dose 30 µM, n = 13; dose 100 µM, n = 11) or BLA-projecting (purple; dose 10 µM, n = 13; dose 30 µM, n = 8; dose 100 µM, n = 10) VTA DA neurons (two-way RM ANOVA: dose effect, F_2,57_ = 9.54, ***p < 0.001; projection effect, F_1,57_ = 16.77, ***p < 0.001). **(C)** Left: example electrophysiological traces of sIPSCs before and after nicotine bath application and associated nicotinic currents. Right: mean sIPSC amplitude and frequency before and after nicotine in either BLA-projecting (n = 14, *top*) or NAcLSh-projecting (n = 17, *bottom*) VTA DA neurons (Welch two sample t-test, t_30.343_ = -2.14, *p = 0.04). **(D)** Right: AAVretro.mCherry.Cre was injected in the NAc (NAcLShell + NAcMShell + NAcCore) together with a Cre-dependent lentivirus for re-expression of the β2 nAChR subunit in the VTA. Left: immunohistochemistry showing viral expression of mCherry and GFP in tyrosine hydroxylase-expressing DA neurons (TH) in the VTA. **(E)** Density of responses evoked by i.v. injections of nicotine (Nic; black) or saline (Sal; gray) in VTA DA neurons from GFP^NAcVec^ (n = 19, *top*) or β2^NAcVec^ mice (n = 32, *bottom*). Responses are expressed as a percentage of the change in firing frequency induced by the injection (Kolmogorov-Smirnov test: comparison between saline and nicotine responses for β2^NAcVec^: D = 0.38, *p = 0.02; comparison of nicotine responses between β2^NAcVec^ and GFP^NAcVec^: D = 0.38, *p = 0.05). **(F)** Left: time course of the mean change in firing frequency (% of baseline) after i.v. nicotine or saline (gray) injection for VTA DA neurons from GFP^NAcVec^ (*top*) or β2^NAcVec^ mice (*bottom*). Right: comparison of firing rate variation from baseline after saline and nicotine injection for nicotine-activated (red) or nicotine-inhibited (blue) VTA DA neurons (n = 17 and 15; Student paired t-test: Bas vs Nic+, t_16_ = -4.46, ***p < 0.001; Bas vs Nic-, t_14_ = 3.46, **p = 0.004; paired Wilcoxon test for comparison between saline-induced and nicotine-induced variations: for Nic+ neurons, V = 0, ***p < 0.001; for Nic-neurons, V = 1, ***p < 0.001). **(G)** Same as (D) but AAVretro.mCherry.Cre was injected in the Amg (BLA + CeA). **(H)** Same as (E) for GFP^AmgVec^ (n = 10, *top*) and β2^AmgVec^ (n = 29, *bottom*; Kolmogorov-Smirnov test, D = 0.45, **p = 0.005). **(I)** Same as (F) for GFP^AmgVec^ (n = 10, *top*) and β2^AmgVec^ (n = 13 and 9, respectively; Student paired t-test: Bas vs Nic+, t_12_ = -6.99, ***p < 0.001; Bas vs Sal-, t_8_ = 7.52, ***p < 0.001; Bas vs Nic-, t_8_ = 5.14, ***p < 0.001; paired Wilcoxon or Student t-test for comparison between saline-induced and nicotine-induced variations: for Nic+ neurons, V = 91, ***p < 0.001; for Nic-neurons, t_8_ = 0.87, p = 0.4).

We then investigated whether nicotine-induced inhibition could be mediated by local GABAergic interneurons. Nicotine (30 µM) was perfused locally and nicotine-evoked GABAergic currents were recorded within the two subpopulations. In Amg-projecting VTA DA neurons, which are known to be inhibited *in vivo*, nicotine had no effect on the amplitude or frequency of spontaneous postsynaptic inhibitory currents (sIPSCs; **Figure 2C, *top***; **Figure S3A**). Conversely, NAc-projecting VTA DA neurons showed an increase in sIPSC amplitude on average (**Figure 2C, *bottom***) and in cumulative distributions after nicotine (**Figure S3A**), despite the fact that these neurons are typically activated by nicotine *in vivo*. Indeed, bath application of nicotine to the slice evoked a long-lasting nicotinic inward current of larger amplitude in NAc-projecting DA neurons (**Figure S3B**), in line with our observations using puff applications of nicotine. In addition, the mean amplitude and frequency of miniature postsynaptic inhibitory currents (mIPSCs) per neuron were similar in the two subpopulations, indicating that there is no major difference in overall GABAergic inputs (**Figure S3C, D**). Taken together, these results suggest that nicotine-induced inhibition of Amg-projecting DA neurons is unlikely to be mediated primarily by an increased GABAergic signaling from local VTA interneurons.

### Nicotine-induced activation of NAc-projecting DA neurons is sufficient to trigger inhibition of VTA DA neurons

Our *ex vivo* investigation revealed that NAc-projecting DA neurons show a stronger activation in response to directly-applied nicotine than Amg-projecting DA neurons. Therefore, we explored whether the inhibition of Amg-projecting DA neurons could result from the activation of NAc-projecting DA neurons. Initially, we tested whether the selective activation of NAc-projecting DA neurons by nicotine could lead to the inhibition of VTA DA neurons *in vivo*. In β2^-/-^ mice, a retrograde virus expressing a Cre recombinase was injected into the three subareas of the NAc (NAcLSh, NAcMSh, NAcCore) and a Cre-dependent lentivirus expressing the β2 subunit was injected into the VTA. This approach restricted the expression of the β2 subunit in NAc-projecting neurons. We then performed *in vivo* juxtacellular recordings of VTA DA neurons in vectorized mice expressing either GFP alone, as control (GFP^NAcVec^), or β2-GFP (β2^NAcVec^; **Figure 2D**). In β2^NAcVec^ mice, i.v. nicotine induced firing variations characterized by a bimodal distribution different from saline, which was not observed in GFP^NAcVec^ control mice (**Figure 2E)**. VTA DA neurons of GFP^NAcVec^ mice did not respond to nicotine, whereas nicotine-induced activation and inhibition were restored in β2^NAcVec^ mice (**Figure 2F**). Following the same strategy, we also selectively re-expressed the β2 subunit in the VTA-Amg pathway of β2^-/-^ mice (β2^AmgVec^; **Figure 2G)**. In β2^AmgVec^ mice, nicotine-induced responses exhibited a unimodal right-shifted distribution compared to those induced by saline, but were not significantly different from those observed in GFP^AmgVec^ mice (**Figure 2H**). Nicotine-activated neurons showed a firing rate variation statistically different from saline injection, but no significant inhibition of DA neurons in response to nicotine was observed in β2^AmgVec^ mice (**Figure 2I)**. Thus, specific nicotine-induced activation of NAc-projecting, but not Amg-projecting DA neurons, was sufficient to restore nicotine-induced inhibition in a subset of DA neurons.

This indicated that nicotine-induced inhibition is a consequence of the activation of VTA-NAc projections. Such a mechanism of inhibition could arise from the activation of D2 autoreceptors following somatodendritic DA release from VTA DA neurons that have been activated by nicotine. To investigate this possibility, we examined *in vivo* responses to nicotine in VTA DA neurons from DAT-D2R^-/-^ mice, which specifically lack D2 receptors in DA neurons. Our findings indicate that both excitatory and inhibitory responses were unaffected in these mice, indicating that D2 autoreceptors do not underlie nicotine-induced inhibition (**Figure S4**).

Overall, these results indicate that nicotine-induced inhibition does not rely on local GABAergic circuitry or DA release. Consequently, we proceeded to test whether it may instead be triggered by the activation of VTA-NAc projections, which in turn engage long-range GABAergic neurons targeting the VTA.

### Amg-projecting DA neurons are targeted by inhibitory inputs from the NAc

NAc medium spiny neurons expressing dopaminergic D1 receptors (D1-MSNs) are known to send direct inhibitory projections to VTA neurons ^20–22^. First, we examined the connectivity between NAc GABAergic terminals in the VTA and BLA-projecting DA neurons, by combining optogenetics with patch-clamp recordings (**Figure 3A**). ChR2 was expressed in the three subareas of the NAc (in different batches of mice), red retrobeads were injected into the BLA, and light-evoked IPSCs were recorded in BLA-projecting DA neurons. Neurons were then filled with biocytin to verify their DA phenotype (**Figure 3B**). We found that a significant percentage of BLA-projecting DA neurons responded to light stimulation from the NAcMSh (41.2%, n=7/17 neurons), while a higher proportion responded to light stimulation from the NAcLSh (61.6%, n=8/1 neurons) and almost all responded to light stimulation from the NAcCore (90.9%, n=10/11 neurons; **Figure 3C**). These data show that VTA DA neurons projecting to the BLA receive strong GABAergic inhibition emanating from the NAc Shell and Core.

**Figure 3:**
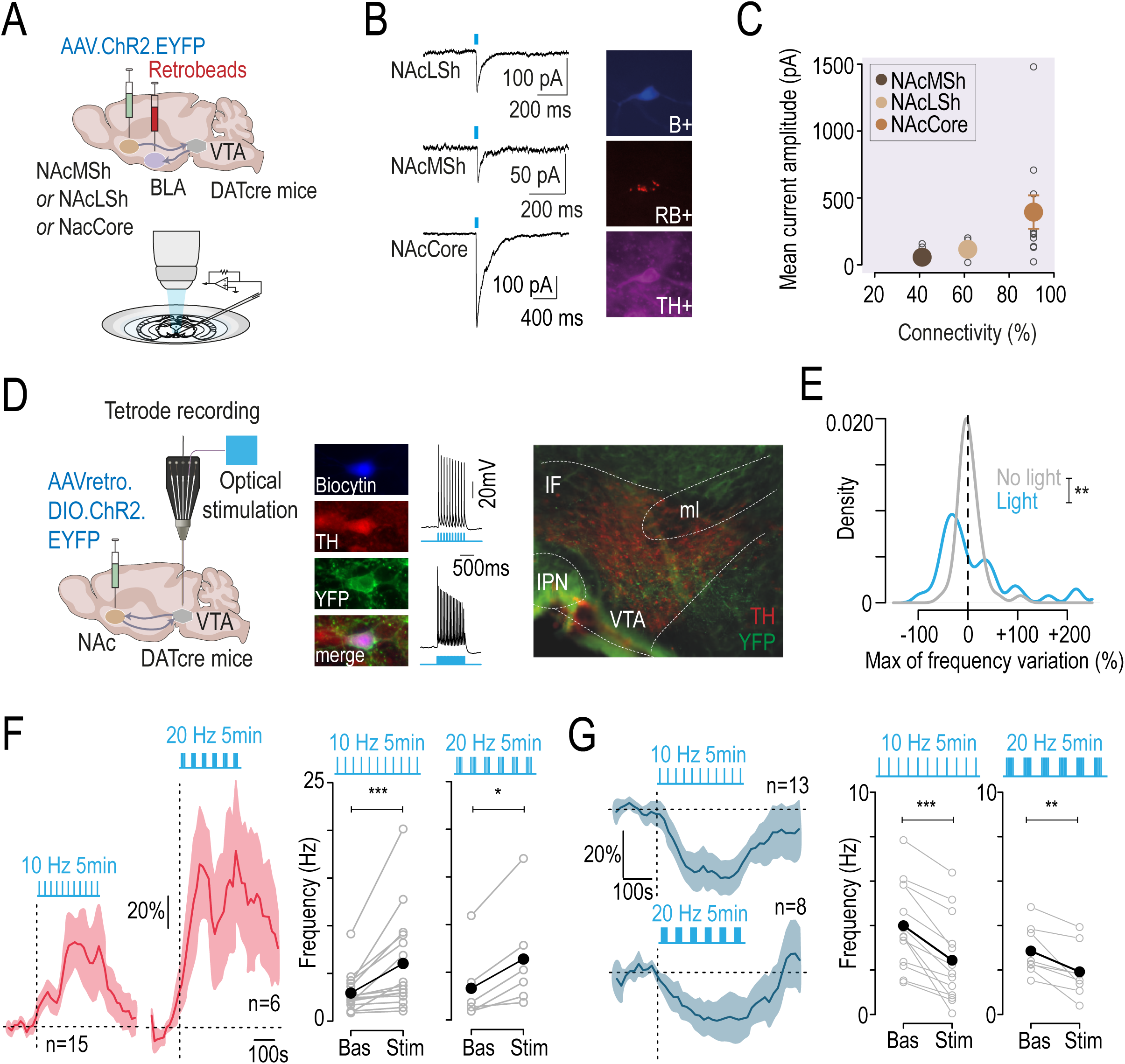
VTA_DA_-NAc pathway activates inhibitory feedback targeting VTA-BLA neurons and inducing inhibition in the VTA. **(A)** Retrobeads were injected in the BLA and ChR2 was injected in the NAc (NAcLShell or NAcMShell or NAcCore). **(B)** Example electrophysiological recordings of light-evoked IPSCs in BLA-projecting DA neurons. Post hoc identification of BLA-projecting DA neurons by immunofluorescent co-labeling of tyrosine hydroxylase (TH), neurobiotin (NB), and retrobeads (RB). **(C)** Synaptic connectivity plotted against the amplitude of light-evoked IPSCs from NAcMSh, NAcLSh or NAcCore projections on BLA-projecting VTA DA neurons (n = 7/17, 8/13 and 10/11 neurons). **(D)** Left: An AAV for Cre-dependent, retrograde expression of ChR2 was injected in the NAc (NAcLShell + NAcMShell + NAcCore) of DAT-Cre mice, and patch-clamp or multi-unit extracellular recordings were performed. Middle: YFP-positive cells recorded in patch-clamp experiments were responsive to 10 and 20 Hz stimulation (ultra-high-power LED: 450-465 nm). Post hoc immunofluorescence verification of co-labeling of TH, YFP and Biocytin. Right: YFP signal was localized to the medial and lateral parts of the VTA. **(E)** Density of responses evoked by light stimulation (ultra-high-power LED: 450-465 nm; 5 min stimulation) at 10 Hz (blue) and spontaneous firing variation during baseline (gray). Light-evoked responses are expressed as a percentage of the change in firing frequency induced by light (n = 43; Kolmogorov-Smirnov test: D = 0.4, **p = 0.002). **(F)** Left: time course of the average change in firing frequency (% of baseline) upon light stimulation at 10 or 20 Hz in light-activated pDA neurons (red). Right: comparison of firing rate variation between baseline and light stimulation for 10 or 20 Hz stimulation pattern (n = 15 and 6, respectively; paired Wilcoxon test: for 10 Hz, V = 0, ***p < 0.001; for 20 Hz, V = 0, *p = 0.03). **(G)** Left: time course of the average change in firing frequency (% of baseline) upon light stimulation at 10 or 20 Hz in light-inhibited pDA neurons (blue). Right: comparison of firing rate variation between baseline and light stimulation for 10 or 20 Hz stimulation pattern (n = 13 and 8, respectively; paired Student t-test: for 10 Hz, t_12_ = 6.1, ***p < 0.001; for 20 Hz, t_7_ = 3.87, **p = 0.006).

### NAc-projecting DA neurons trigger inhibitory feedback on VTA DA neurons

In light of these results, we sought to test whether activation of the VTA-NAc dopaminergic pathway (VTA_DA_-NAc) leads to inhibition of VTA neurons by recruiting descending GABAergic inputs from the NAc. To investigate the functional impact of this pathway, we injected a retrograde ChR2-containing virus (retroChR2) with YFP signal into the three subareas of the NAc of DATcre mice, for restricted expression of ChR2 in VTA DA neurons projecting to the NAc (**Figure 3D**, ***left***). All YFP-positive cells recorded in patch-clamp experiments followed 10 and 20 Hz stimulation patterns (**Figure 3D**, ***middle***). Recorded neurons were confirmed as DA by post-hoc immunofluorescence verification (TH+, YFP+, B+). Neurons expressing the YFP signal were localized in the medial and lateral parts of the VTA (**Figure 3D**, ***right***), consistent with previous findings^8,22^.

We then perfomed multi-unit extracellular recordings combined with optical stimulation in the VTA. The optical fiber was positioned laterally within the VTA, to maximize photoactivation of NAc-projecting DA neurons, which were found to be predominantly lateralized (**Figure S2**,^16^). To mimic nicotine-induced excitation, we applied 5 min of optogenetic stimulation at 10 or 20 Hz, delivered in a regular or burst pattern, respectively. This resulted in frequency variations in putative DA (pDA) neurons, revealing a bimodal distribution of the frequency variation that differed from the maximum spontaneous variation occurring during baseline (n = 43, **Figure 3E**). Specifically, when exposed to 10 or 20 Hz photostimulation, a significant increase in firing frequency was observed in a subset of pDA neurons, likely those that express ChR2 (**Figure 3F**). Simultaneously, a significant proportion of pDA neurons were inhibited by the photostimulation (**Figure 3G**). As we are unable to label neurons *in vivo* using this electrophysiological technique, and therefore unable to confirm their projection site, we tested the response to nicotine in 6 light-responding neurons. We found that 4 out of the 5 neurons inhibited by 10 Hz photostimulation were also inhibited by nicotine (**Figure S5A**), suggesting that some of the light-inhibited neurons may project to the Amg. These results provide compelling evidence that activation of NAc-projecting DA neurons, independent of nicotine, triggers inhibition of another subset of VTA DA neurons.

Taken together, these results suggest that activation of NAc-projecting DA neurons leads to subsequent activation of MSNs in the NAc. Since these MSNs have established connections with BLA-projecting VTA DA neurons, they may be involved in a VTA_DA_-NAc feedback inhibitory loop. This loop, triggered by nicotine or ethanol, may explain the observed inhibitory effects on a specific DA subpopulation.

### Inhibitory feedback from the NAc mediates nicotine-induced inhibition of DA neurons

To confirm that nicotine-induced inhibition involves descending fibers from the NAc to the VTA, we expressed the halorodopsin JAWS in the three subregions of the NAc of WT mice, to block any NAc feedback onto the VTA (**Figure 4A**). We then recorded the response of VTA pDA neurons to an i.v. injection of nicotine, focusing on neurons inhibited by the drug, while simultaneously photoinhibiting NAc terminals in the VTA. We found that in most cases, photoinhibition reverse nicotine-induced inhibition (**Figure 4B, C, *left***). Specifically, compared to saline, maximum variations in firing rate from baseline showed a significant inhibition when nicotine was injected alone, but not when combined with photoinhibition of NAc terminals, where 5 out of 6 neurons showed an increase in firing rate (**Figure 4C, *right***; **Figure S5C**). Photoinhibition alone had no effect on firing frequency, suggesting that the change in nicotine’s effect during light is not due to disinhibition (**Figure S5C**). Similar experiments on nicotine-activated neurons reveal that photoinhibition of NAc terminals did not prevent nicotine-induced activation of DA neurons (**Figure S5D, E**). Thus, photoinhibition of NAc terminals in the VTA prevents nicotine-induced inhibition of DA neurons by blocking inhibitory feedback.

**Figure 4:**
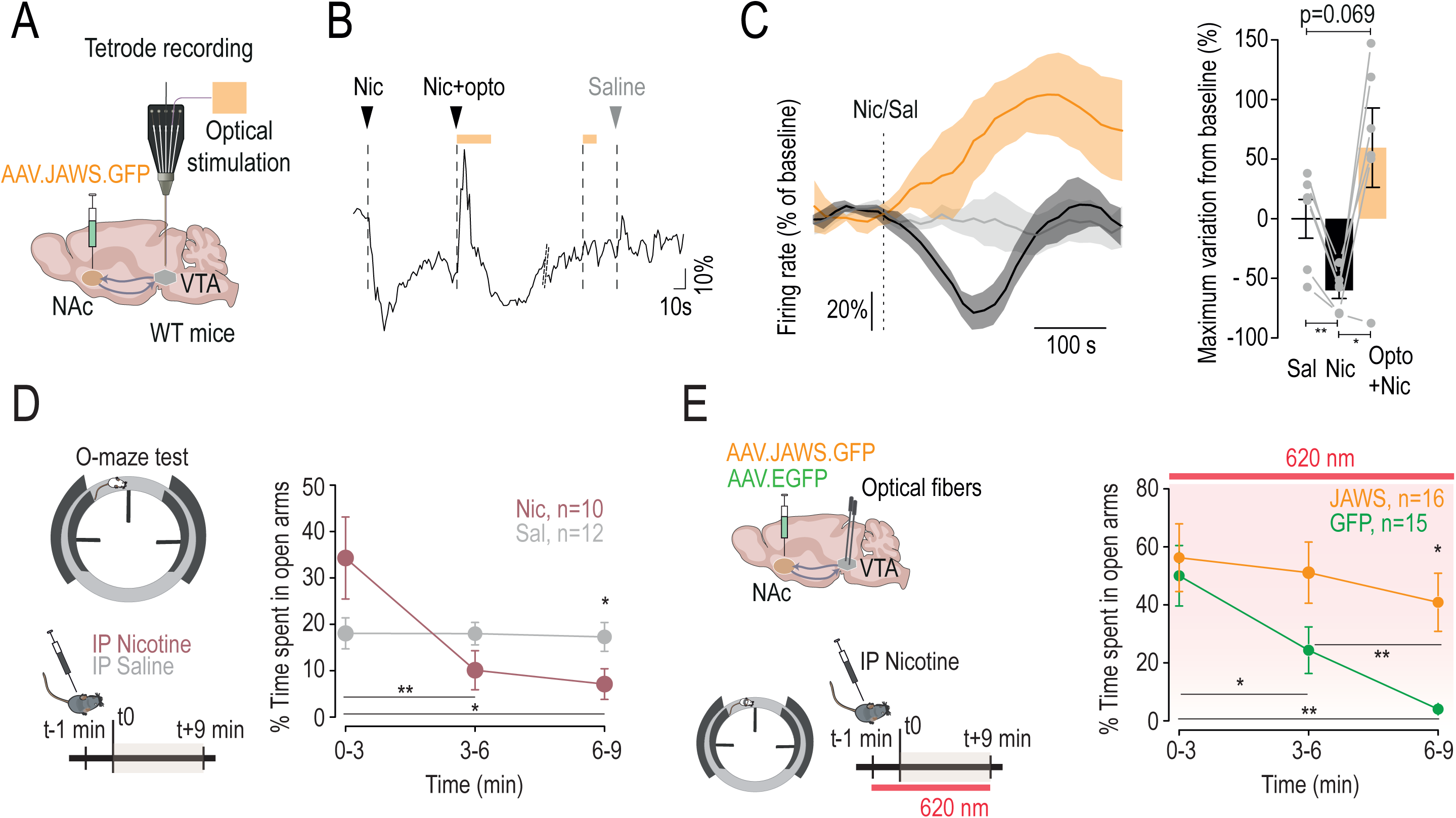
Inhibitory feedback loop from the NAc mediates nicotine-induced inhibition of VTA pDA neurons and nicotine-induced anxiety. **(A)** A JAWS-expressing AAV was injected in the NAc (NAcLShell + NAcMShell + NAcCore) and multi-unit extracellular recordings were performed. **(B)** Example of change in firing rate variation (% of baseline) after i.v. nicotine injection, i.v. nicotine injection coupled with optogenetic inhibition of NAc terminals in the VTA (5 min, continuous stimulation), optogenetic inhibition of NAc terminals alone, or i.v. saline injection. **(C)** Left: time course of the mean change in firing frequency (% of baseline) after i.v. nicotine of pDA inhibited neurons (n = 6) with (orange) or without (black) optogenetic inhibition of NAc terminals in the VTA. Right: comparison of maximum change from baseline between i.v. saline injection, i.v. nicotine injection alone, or i.v. nicotine injection + optogenetic inhibition of NAc terminals in the VTA (pairwise comparisons using paired t-test: Sal vs Nic, t_5_ = 5.76, p = 0.007; Nic vs Nic + Opto, t_5_ = -3.98, **p=0.02; Sal vs Nic + Opto, t_5_ = -2.31, p = 0.07). **(D)** Left: intraveneous (i.p.) injection of nicotine (0.5 mg/kg; n = 10) or saline (n = 12) 1 min before the elevated O maze (EOM) test. Right: percentage of the time spent in open arms (two-way RM ANOVA: group x time interaction, F_2,40_ = 8.66, ***p < 0.001; main effect of time, F_2,40_ = 7.81, **p = 0.001; post-hoc pairwise comparisons using Wilcoxon with Holm corrections for time effect on mice having received i.p. Nic: for 3 vs 6min, V = 55, **p = 0.006; for 3 vs 9min, V = 51, *p = 0.03; for 6 vs 9min, V = 22.5, p = 0.57; post-hoc Wilcoxon test at 9min between Sal and Nic, W = 94, *p = 0.03). **(E)** Left: injection of JAWS- or GFP-expressing AAV in the NAc (NAcLShell + NAcMShell + NAcCore) and implantation of optical fibers in the VTA. I.p. nicotine (0.5 mg/kg) was performed 1 min before the EOM test and light stimulation was then continuously maintained throughout the entire test. Right: percentage of the time spent in open arms with continuous inhibition of NAc terminals in the VTA (two-way RM ANOVA: group x time interaction, F_2,58_ = 3.27, *p = 0.04; group effect, F_1,29_ = 4.18, *p = 0.05; time effect, F_2,58_ = 12.26; ***p < 0.001; post-hoc pairwise comparisons using Wilcoxon with Holm corrections for GFP mice: for 3 vs 6min, V = 85, *p = 0.04; for 3 vs 9min, V = 89, **p = 0.006; for 6 vs 9min, V = 90, p = 0.006; post-hoc Wilcoxon test at 9min between JAWS and GFP: W = 58, *p = 0.01).

### Inhibitory feedback from the NAc mediates nicotine-induced anxiety-like behavior

We next investigated the implication of this NAc-VTA inhibitory feedback in the anxiogenic effect of nicotine. Intraperitoneal (i.p) injection of nicotine (0.5 mg/kg) induced a decrease in the time spent in open arms in an elevated O maze (EOM), indicative of an anxiogenic effect of the drug (**Figure 4D**), which was not correlated with locomotor activity over time (**Figure S6A**). We have previously shown that this anxiogenic effect of nicotine requires the inhibition of VTA DA neurons projecting to the Amg^16^. To complete our demonstration of a functional VTA-NAc inhibitory loop, we examined whether blocking the inhibitory feedback originating from the NAc could alleviate the anxiogenic effect of nicotine. To this aim, we expressed JAWS or GFP in the three subareas of the NAc in WT mice and implanted optical fibers bilaterally in the VTA (**Figure S6B**). Mice then received an i.p. injection of nicotine (0.5 mg/kg) 1 min before the EOM test and light stimulation in the VTA throughout the 9 min test. Photoinhibition of NAc terminals in the VTA during the EOM test abolished the anxiogenic effect of nicotine injection, as indicated by the percentage of time spent by JAWS-expressing mice in the EOM open arms that did not decrease during the test, in contrast to GFP-expressing mice (**Figure 4E**). These effects were not correlated with locomotor activity over time (**Figure S6C**). Without nicotine, photoinhibition of NAc-VTA terminals alone in JAWS-expressing mice had no effect on the distance travelled in the open field or the time spent in the open arms of the EOM test (**Figure S6D, E**). Finally, considering that both nicotine and ethanol inhibit Amg-projecting DA neurons, we examined whether ethanol, like nicotine, could induce changes in anxiety levels in mice through the NAc-VTA circuit. As observed with nicotine, inhibition of NAc terminals prevents the decrease in time spent in the open arms of the EOM over time following ethanol injection (**Figure S6F**), suggesting that the same inhibitory feedback loop is involved in these effects of ethanol. These data highlight the functional role of an inhibitory loop between the VTA and the NAc in mediating nicotine-induced anxiety-like behavior, primarily through the inhibition of DA neurons.

## DISCUSSION

The notion that drugs reinforce behavior through a broad activation of the DA system is challenged by evidence showing distinct subpopulations of DA neurons in the VTA, each associated with specific appetitive, aversive, or attentional behaviors^7,8,23,24^. Our previous work has highlighted the heterogeneity of the VTA, particularly in response to nicotine^15,16^. Nicotine triggers both activation and inhibition of VTA DA neurons, associated with two distinct anatomical and functional circuits. These circuits produce opposite behavioral effects: inhibition of VTA DA neurons projecting to the BLA is anxiogenic, whereas activation of those projecting to the NAcLSh is rewarding^16^. During nicotine exposure, these dual DA pathways may act concurrently, as the same nicotine dose can be both rewarding and anxiogenic^16,25^. Here, we present evidence that inhibition and activation are not isolated events; rather, inhibition arises as a consequence of activation and is mediated by feedback inhibition from the NAc. This finding has significant conceptual implications, suggesting that the VTA should not be viewed as a collection of independent subcircuits that function in isolation. Instead, it suggests a more integrated and interconnected system where activation in one pathway can directly influence activity in another.

In this study, we propose that nicotine-induced inhibition of DA neurons arises from two key properties of the VTA-NAc network: the recurrent architecture of this network and a differential sensitivity to nicotine of DA neurons projecting to the Amg or NAc. Recent studies have shown that VTA DA neurons form organized reciprocal connections with the NAc subregions they target, establishing distinct inhibitory feedback loops^22^. Our findings demonstrate that Amg-projecting DA neurons receive significant inhibitory inputs from all NAc subregions, and that optogenetic inhibition of these inputs prevents nicotine-induced inhibition of VTA DA neurons. This indicates that activation of NAc-projecting DA neurons may trigger feedback inhibition in the VTA through NAc-VTA projections that particularly target Amg-projecting neurons. Both activation and inhibition of DA neurons in response to nicotine are mediated by β2*nAChRs in the VTA^16^. Nicotine binds to nAChRs expressed by DA and GABA neurons in the VTA^26^, directly exciting DA neurons and indirectly inhibiting them through activation of GABA neurons^14^. Unexpectedly, nicotine increases sIPSCs in NAc-projecting DA neurons but not in Amg-projecting ones, indicating preferential activation of GABAergic interneurons targeting NAc-projecting neurons. Despite this GABAergic influence, we found stronger nicotine-evoked depolarizing currents in NAc-projecting neurons, which aligns with the increased firing rates in response to nicotine *in vivo*. These observations are consistent with previous results demonstrating the necessity of co-activating DA neurons and GABAergic interneurons by nicotine for DA release in the NAc^14^. In contrast, weaker nicotinic currents in Amg-projecting DA neurons suggest that the direct excitatory effect of nicotine is not sufficient to overcome the inhibition from the NAc, leading to decreased activity in these neurons. Heterogeneity in nicotinic currents among DA neurons, due to differential expression of β2, β4, or α7 nAChRs, has been described, with stronger currents linked to β2*nAChRs^27^. Our findings suggest that β2*nAChRs are more abundantly expressed in NAc-projecting DA neurons, resulting in their preferential activation over Amg-projecting neurons when nicotine binds to VTA nAChRs *in vivo*. Using retrograde viral rescue to restrict β2*nAChRs expression to NAc-projecting neurons, we demonstrated that nicotine activation of these neurons can consequently inhibit other VTA DA neurons, presumably those lacking β2*nAChRs. Consistent with studies showing that intra-VTA nicotine infusion increases the firing of MSNs^28^, our results suggest that nicotine efficiently triggers inhibitory feedback from the NAc to the VTA via NAc-projecting DA neurons. However, our viral rescue strategy has limitations, such as the inability to specifically target DA neurons and potential discrepancies in the re-expression levels of β2*nAChRs. In addition, other nAChR subtypes in the NAc or regions targeting the VTA may also be involved. Optogenetic activation of the VTA_DA_-NAc pathway in DAT-Cre mice replicated the nicotine-induced activation/inhibition pattern, confirming that NAc-projecting DA neurons can trigger inhibition in VTA DA neurons. Optogenetic-induced inhibition of VTA neurons correlates with nicotine-induced inhibition, further suggesting that activation of the VTA_DA-_ NAc pathway is a key component of the DA inhibition we observed.

Here, we provide evidence that nicotine-induced activation of the VTA_DA_-NAc pathway leads to inhibition of the VTA_DA_-Amg pathway, highlighting a complex interplay between reward-related and emotion-related neural circuits during drug exposure. Nicotine exhibits both rewarding^3,14^ and anxiety-inducing properties^29–31^ that are linked to DA signaling. Inhibition of the VTA_DA_-Amg pathway increases anxiety, whereas its activation decreases it^16^. Conversely, activation of the VTA_DA_-NAc pathway triggers online conditioned place preference, which is associated with reward signaling. Furthermore, counteracting the inhibition of BLA DA terminals during nicotine exposure abolishes nicotine-induced anxiety, while counteracting the activation of NAc lateral shell DA terminals reduces it^16^. Consistent with these findings, we abolished nicotine-induced anxiety associated with inhibition of Amg-projecting DA neurons by inhibiting NAc terminals in the VTA. Therefore, NAc-VTA projections targeting Amg-projecting DA neurons may link the rewarding and negative emotional outcomes of nicotine intake. Recent studies have investigated the effect of optogenetic modulation of NAc-VTA terminals originating from discrete parts of NAc subregions in relation to anxiety. These studies show no effect on anxiety with activation^22^ and an anxiogenic-like effect with inhibition^32^. This may seem to contradict our findings showing that activation of NAc-VTA projections is necessary for nicotine-induced anxiety-like behavior. However, it is important to note that discrete optogenetic modulation, while effective for precise functional dissection of neural circuits, cannot recapitulate the simultaneous and widespread activation of networks that occurs during drug exposure. Indeed, nicotine indiscriminately activates DA neurons projecting to different subregions of the NAc^16^, which in turn send inhibitory inputs to Amg-projecting DA neurons. Our approach of globally activating VTA_DA_-NAc pathways or inhibiting NAc-VTA projections aims to mimic or suppress nicotine effects, respectively. Our results suggest that inhibition of Amg-projecting DA neurons and the associated anxiety-like behavior require significant recruitment of the VTA-NAc inhibitory projections. Furthermore, while the NAc-VTA projections appear critically involved in nicotine-induced anxiety, other pathways likely also contribute to the anxiogenic effects of nicotine.

An important aspect of our findings is that alcohol, like nicotine, induces inhibition of Amg-projecting neurons and activation of NAc-projecting neurons. This correlation suggests that both substances affect overlapping neural circuits within the VTA. Although alcohol and nicotine rely on different molecular targets - nicotine through nAChRs and alcohol through other mechanisms^18^ - both activate NAc-projecting neurons and increase DA release in the NAc^1^.This suggests that alcohol may also elicit inhibitory feedback from the NAc, which is consistent with our results showing that inhibition of NAc-VTA projections interferes with the effect of ethanol on anxiety levels. Unlike nicotine, there is no evidence of a difference in alcohol sensitivity in a specific DA subpopulation. However, recent research shows greater activation of DA neurons in the lateral VTA that project to the NAc in response to ethanol, compared to medially located neurons^17^.This hints that ethanol, like nicotine, may override potential inhibitory mechanisms in a specific DA subpopulation. Combined with optogenetic activation of NAc-projecting DA neurons, our findings with drugs support that activating the VTA_DA_-NAc pathway can reliably inhibit the VTA_DA_-Amg pathway. In contrast to nicotine and alcohol, natural rewards activate both DA pathways similarly^33^, and with briefer and weaker activation, which may not be sufficient to trigger NAc-VTA inhibitory projections. In addition, inputs mediating natural rewards may activate both NAc- and Amg-projecting neurons, thus overriding any feedback inhibition. Therefore, activation of the VTA-NAc inhibitory feedback loop, leading to opposing DA signaling in reward and emotional circuits, may be characteristic of drugs of abuse. However, drugs that increase DA by acting directly on terminals, such as cocaine and amphetamine, may bypass the functional consequences of this loop, resulting in different rewarding or negative outcomes compared to nicotine or alcohol.

Our study supports the notion that the rewarding signaling of nicotine simultaneously triggers negative emotional states. We demonstrate that acute nicotine induces anxiety as a direct consequence of the activation of the DA reward pathway that supports reinforcement. Thus, anxiogenic and rewarding signals not only occur simultaneously upon nicotine exposure but are also inherently interconnected. This interdependence suggests that alterations in the VTA_DA_-NAc pathway may significantly influence the effects of nicotine on the VTA_DA_-Amg DA pathway, thereby impacting the overall reinforcing properties of the drug. It also suggests that the circuitry underlying negative emotional states, often referred to as the anti-reward system^34^, and the reward circuit evolve in tandem during chronic nicotine exposure and potentially during withdrawal. This raises intriguing question about how two circuits with opposing messages compete to produce nicotine reinforcement, and whether an imbalance between the two could contribute to addiction. The insights provided by our study highlight the intricate neural mechanisms underlying nicotine addiction, particularly the interplay between reward and emotional circuits. Understanding these interactions could pave the way for more effective treatments for nicotine addiction and other substance use disorders where both reward and emotional regulation play crucial roles.

## AUTHORS CONTRIBUTIONS

T.L.B., P.F. and F.M. designed the study. T.L.B., P.F. and F.M. analyzed the data. T.L.B., C.N. and F.M., performed and analyzed *in vivo* juxtacellular electrophysiological recordings. T.L.B. designed, performed and analyzed *ex vivo* patch clamp recordings. E.V. contributed to *ex vivo* patch clamp recordings. T.L.B. performed stereotaxic injections (with contribution from C.S.), fiber implantations and behavioral optogenetic experiments (with contribution from E.V. and L.D.). T.L.B., E.V., L.D., A.G. performed PFA mice intracardiac perfusion and immunostaining experiments. J.J. and N.G performed recordings and signal treatment for *in vivo* multi-unit experiments. S.T. contributed to *in vivo* juxtacellular electrophysiological recordings. C.S., L.M.R. and A.M. contributed to design behavioral optogenetic experiments. S.P., and U.M. provided viruses. E.VJ. provided DAT-D2-knockout (KO) mice. U.M. provided ACNB2-knockout (KO) mice. T.L.B., P.F. and F.M. wrote the manuscript.

## ACKNOWLEDGMENTS

We are grateful to Otilia de Oliveira and Emilie Tubeuf for animal caretaking support. This work was supported by Centre National de la Recherche Scientifique (CNRS; UMR8249), Agence Nationale de la Recherche (Vampire ANR-19-CE16-0001 to F.M.), the French National Cancer Institute Grant (TABAC-16-01, TABAC-19-020 and SPA-21-002 to P.F.), the Foundation for Medical Research (FRM, Equipe FRM DEQ2013326488 to P.F.), the Labex memolife (to J.J. and E.V.), Swedish Research Council (2022-06168 to N.G.), NIDA–Inserm Postdoctoral Drug Abuse Research Fellowship (to L.M.R).

## FIGURES AND LEGENDS

**Figure S1:**
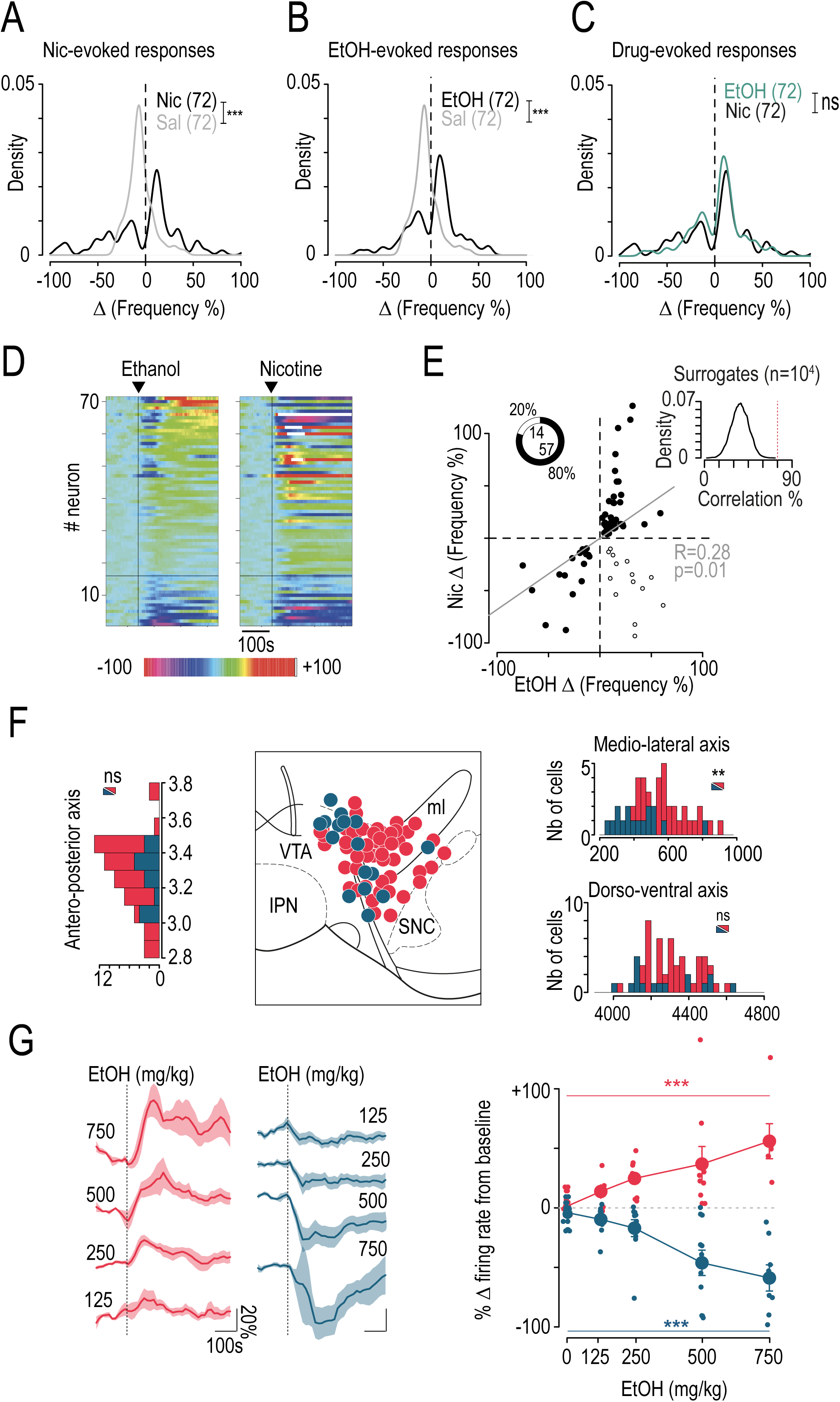
*In vivo* juxtacellular recordings of VTA DA neurons in response to nicotine and ethanol. *Related to* *Figure 1*. **(A)** Density of responses evoked by i.v. injection of nicotine (Nic; black, n = 72) or saline (Sal; gray, n = 72) expressed as a percentage of the change in firing frequency induced by the injection (Kolmogorov-Smirnov test, D = 0.44, ***p < 0.001). **(B)** Density of responses evoked by i.v. injection of ethanol (EtOH; black, n = 72) or saline (Sal; gray, n = 72) expressed as a percentage of the change in firing frequency induced by the injection (Kolmogorov-Smirnov test, D = 0.46, ***p < 0.001). **(C)** Density of responses evoked by i.v. injection of nicotine (Nic; black, n=72) or ethanol (EtOH; green, n=72) expressed as a percentage of the change in firing frequency induced by the injection (Kolmogorov-Smirnov test, D=0.14, p=0.49). **(D)** Individual responses of all VTA DA neurons to ethanol (*left*) and nicotine (*right*) i.v. injections. Neurons are ranked based on their responses to ethanol, from most inhibited (pink) to most activated (white/red). The horizontal line delimits neurons showing inhibition from those showing activation in response to ethanol. **(E)** Correlation between ethanol- and nicotine-induced responses based on the percentage of frequency variation induced by the drugs (Pearson’s correlation, t_69_ = 2.45, R = 0.28, p = 0.02). Percentage and number of correlated (firing rate decrease or increase for both drugs; black dot; n = 57) and uncorrelated (firing rate increase for one drug but decrease for the other; white dot; n = 14) response to nicotine and ethanol. The insert shows the density of the percentage correlation obtained on 10000 surrogates. The red line represents the percentage correlation obtained with the experimental data. **(F)** Localization of DA neurons activated (EtOH+; red; n = 52) and inhibited (EtOH-; blue; n = 15) by i.v. injection of ethanol, positioned on a Paxinos atlas slice at 3.3 mm from bregma, from neurobiotin-filled cell bodies of all recorded neurons. EtOH-neurons had a more medial distribution within the VTA than EtOH+ neurons (Wilcoxon test, **p = 0.003), but neither anteroposterior (Wilcoxon test, p = 0.59) nor dorsoventral (Wilcoxon test, p = 0.24) differences in their distribution were observed. **(G)** Left: time course of mean change in firing frequency (% of baseline) after i.v. injection of different doses of ethanol for activated (EtOH+; red) and inhibited (EtOH-; blue) VTA DA neurons (125, 250, 500 and 750 mg/kg; n = 30/39, 6/10, 9/10, 9/11 and 6/8, respectively for EtOH+ and EtOH-DA neurons). Right: dose-response curves in EtOH+ (red) and EtOH-(blue) VTA DA neurons. Responses to different doses of ethanol are expressed as percentage of variation from baseline (one-way ANOVA: dose effect F_4,55_ = 10.83, ***p < 0.001 and F_4,73_ = 20.64, ***p < 0.001, respectively for EtOH+ and EtOH-DA neurons).

**Figure S2:**
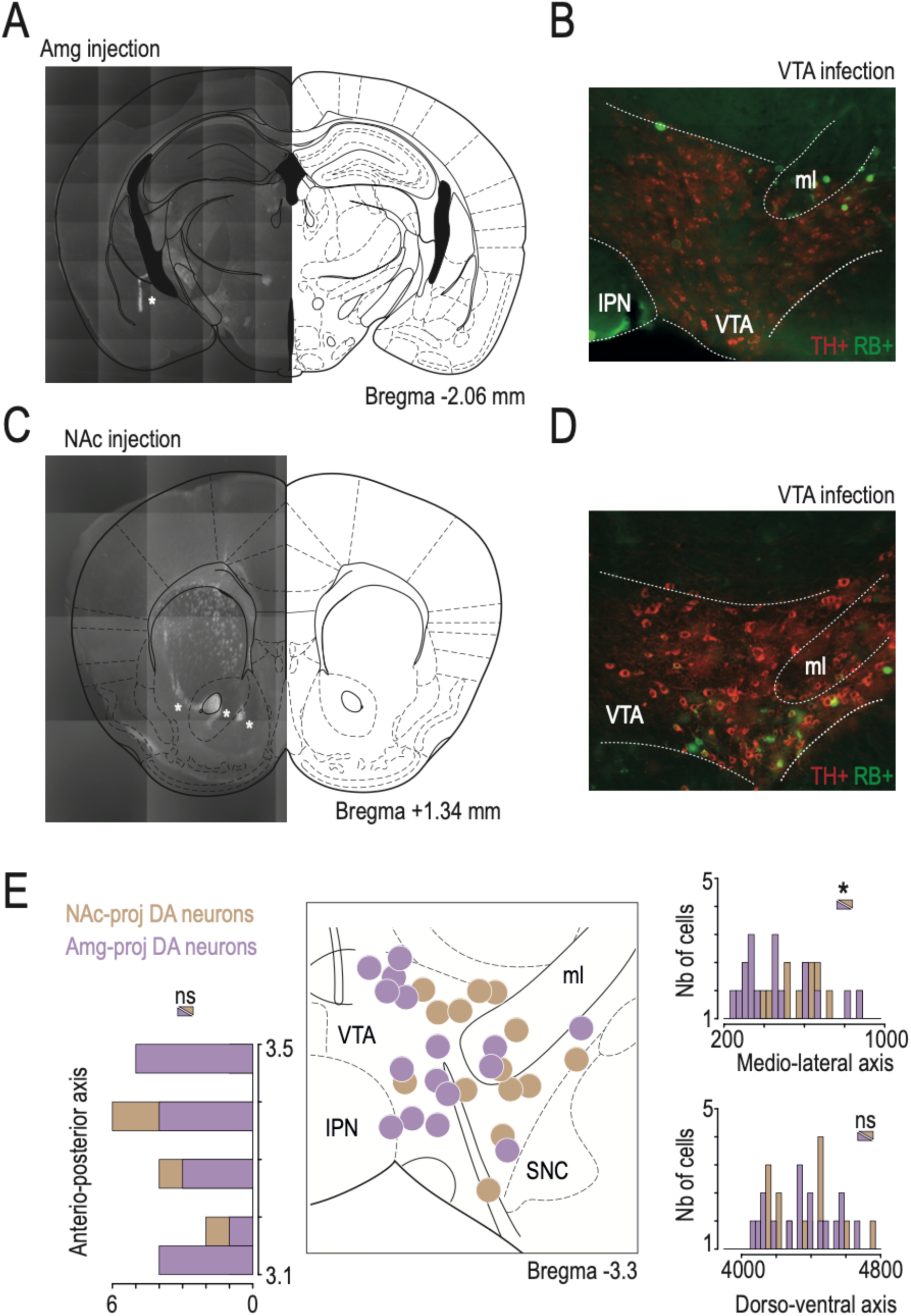
Identification of NAc- and Amg-projecting DA neurons. *Related to* *Figure 1*. **(A)** Example of retrobead (RB) injection sites, indicated by asterisks, in the amygdala (Amg), mapped onto a Paxinos atlas slice. **(B)** Representative immunofluorescence image of VTA slices (TH+, red) revealing neurons containing RB (RB+, green) after RB injection in the Amg. **(C-D)** Same as (A-B) but for RB injection in the nucleus accumbens (NAc). **(E)** Localization of Amg-projecting (purple; n = 16) and NAc-projecting (brown; n = 13) VTA DA neurons, positioned on a Paxinos atlas slice at 3.3 mm from bregma, from neurobiotin-filled cell bodies of all recorded neurons. Amg-projecting DA neurons had a more medial distribution within the VTA than NAc-projecting DA neurons (Wilcoxon test, W=173, *p=0.03), but neither anteroposterior (Wilcoxon test, W=111.5, p=0.77) nor dorsoventral (Welch two sample t-test, t_28.159_ = 0.26, p=0.79) differences in their distribution were observed.

**Figure S3:**
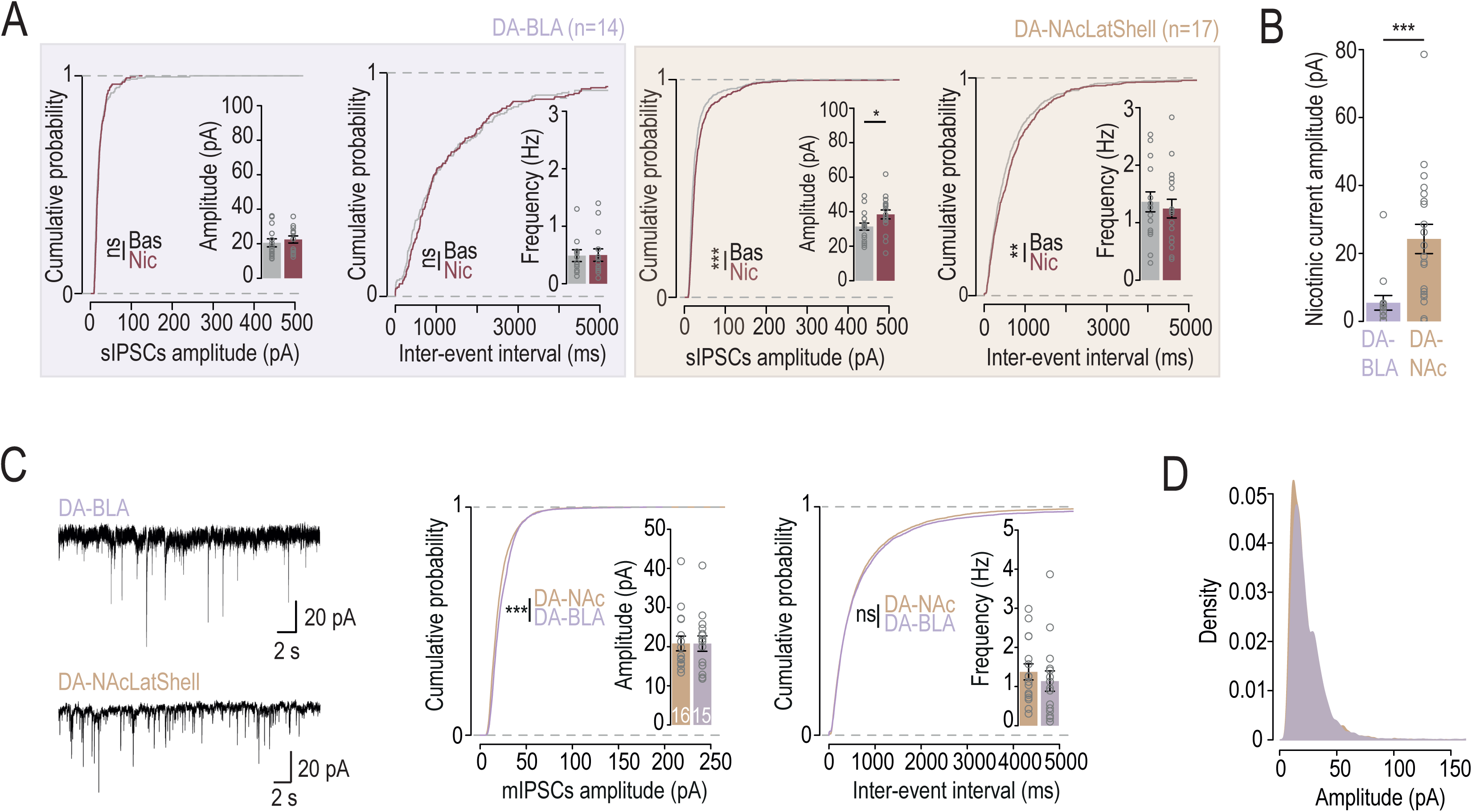
Patch-clamp recordings of spontaneous and miniature inhibitory post-synaptic currents in VTA DA neurons. *Related to* *Figure 2*. **(A)** Cumulative probability and mean plots of sIPSC amplitudes and frequencies, before (gray) and after (red) bath application of nicotine (30 µM), from VTA DA neurons projecting to the NAcLSh (brown) or to the BLA (purple; Kolmogorov-Smirnov test for distributions; NAc-proj sIPSC amplitude: D=0.16, ***p<0.001; NAc-proj sIPSC frequency: D=0.08, p=0.02; Welch two sample t-test for mean plots; NAc-proj sIPSC amplitude, t_30.343_ = -2.135, p=0.04). **(B)** Mean amplitude of nicotinic inward currents evoked by bath application of nicotine (30 µM), concurrent with sIPSCs, in either NAcLSh-(brown; n=17) or BLA-projecting (purple, n=14) VTA DA neurons (Wilcoxon test, W=28, ***p<0.001). **(C)** Example electrophysiological traces of mIPSCs from NAc- or BLA-projecting VTA DA neurons. Cumulative probability and mean plots of mIPSC amplitudes and frequencies from the two populations are represented (Kolmogorov-Smirnov test for amplitudes: D=0.09, ***p<0.001). **(D)** Density plots of mIPSC amplitudes from DA-NAc or DA-BLA VTA neurons.

**Figure S4:**
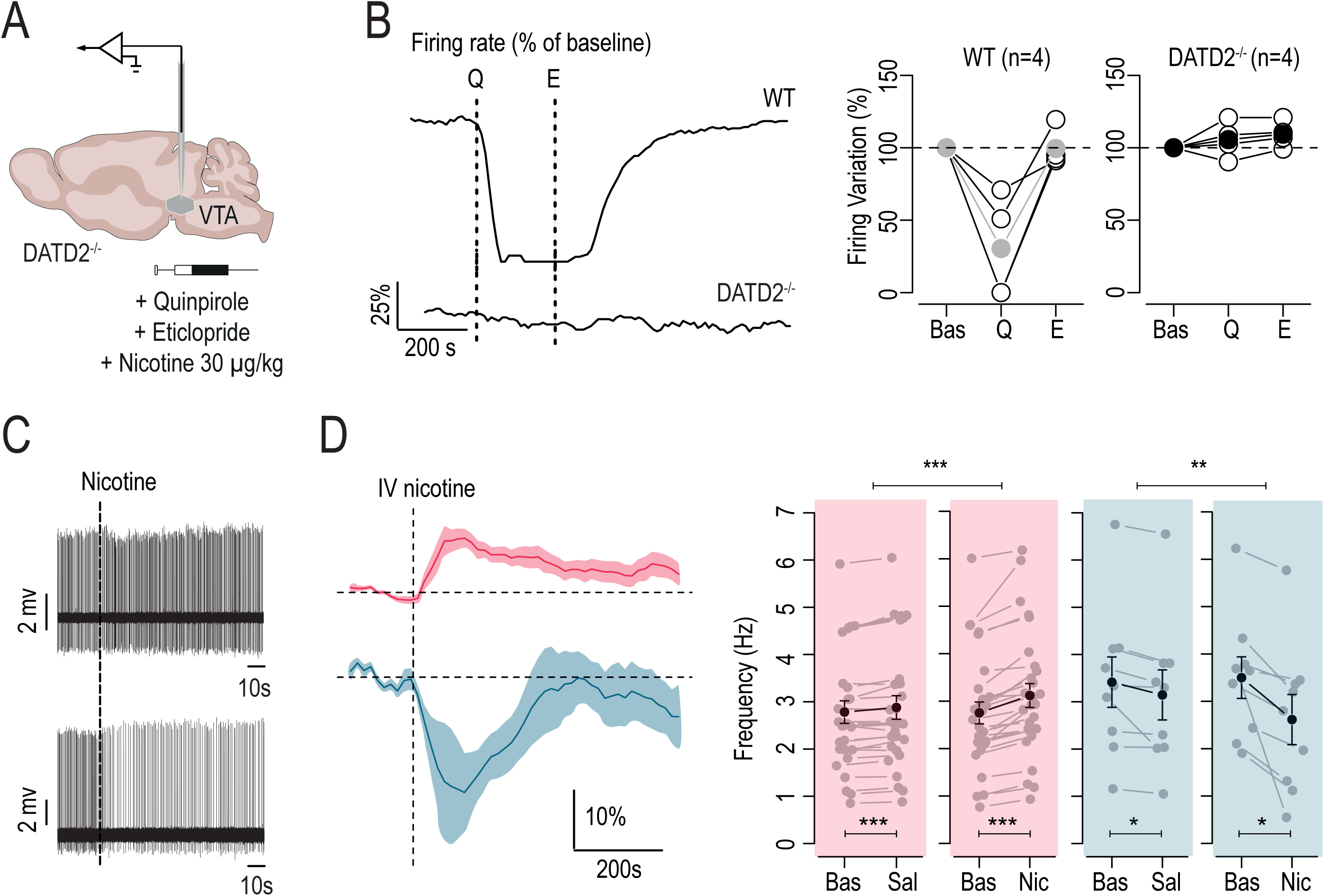
Nicotine-induced responses in DATD2^-/-^ mice. *Related to* *Figure 2*. **(A)** *In vivo* juxtacellular recordings were performed on DATD2^-/-^ mice with i.v. injections of quinpirole (1mg/kg), eticlopride (1mg/kg) or nicotine (30 μg/kg). **(B)** Example and quantification of change in firing rate variation, expressed as percentage of baseline, after i.v. quinpirole and eticlopride injection, on wild-type (WT; n = 4) or DATD2^-/-^ mice (n = 4). **(C)** Representative recordings of VTA DA neurons activated (*top*) or inhibited (*bottom*) by i.v. injection of nicotine. **(D)** Left: time course of the mean change in firing frequency, expressed as percentage of baseline, after nicotine i.v. injection in activated (Nic+, n = 28; red) and inhibited (Nic-, n = 9; blue) VTA DA neurons. Right: comparison of firing rate variation (Hz) between baseline (Bas) and saline (Sal) or nicotine (Nic) injection, in Nic+ and Nic-DA neurons. Maximum firing rate after i.v. nicotine for Nic+ neurons or minimum firing rate after i.v. nicotine for Nic-neurons were represented (paired Student’s t-test: between Bas vs Sal+, t_27_ = -4.73, ***p<0.001, Bas vs Nic+, t_27_ = -6.82, ***p < 0.001, Bas vs Sal-, t_8_ = 2.5, *p = 0.04, Bas vs Nic-, t_8_ = 2.93, *p = 0.02; paired Wilcoxon test: between Bas-Sal vs Bas-Nic, V = 31, ***p > 0.001 and V = 43, **p = 0.01 respectively for Nic+ and Nic-neurons).

**Figure S5:**
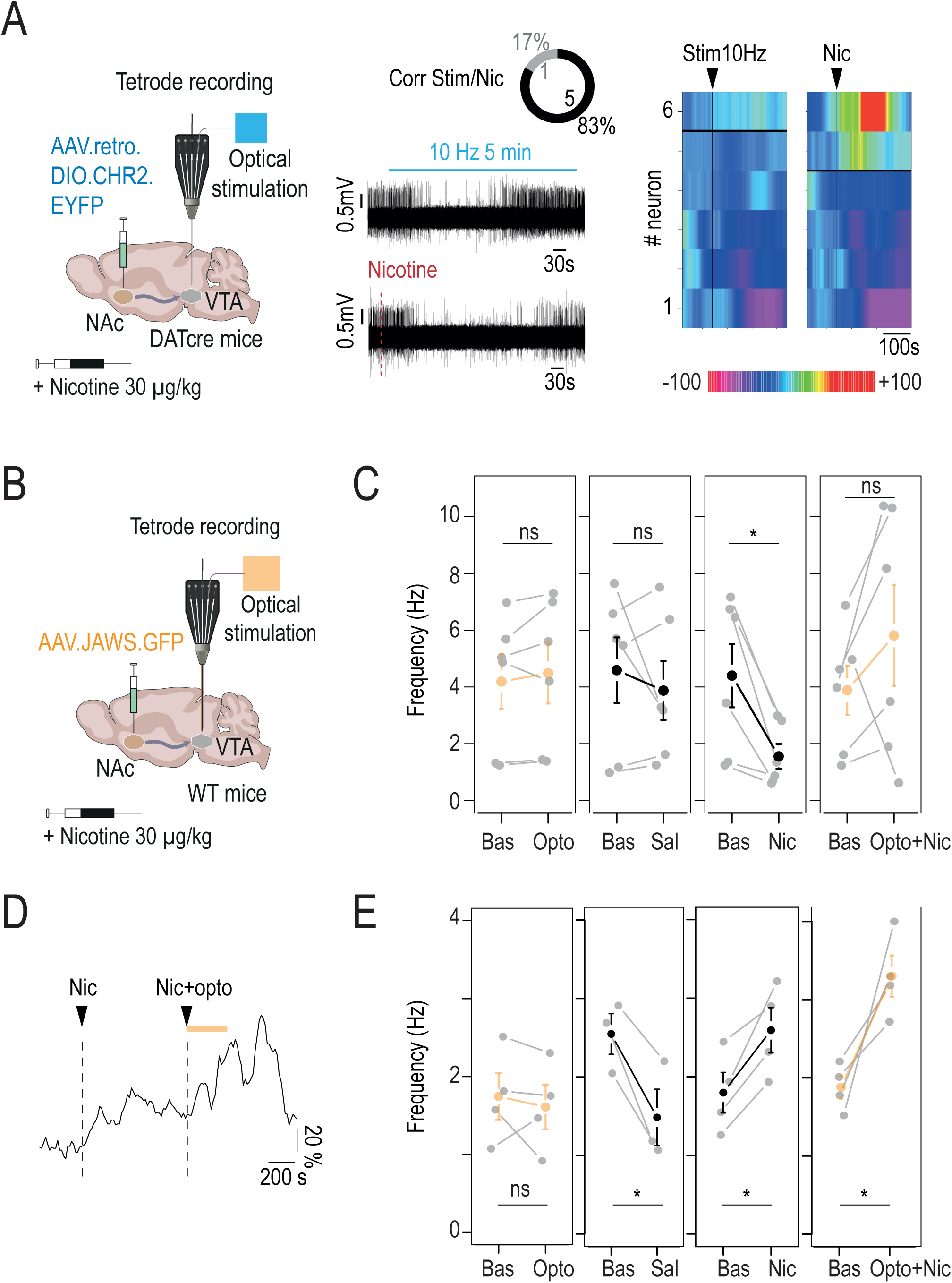
Multiunit recordings of VTA pDA neurons under nicotine and optogenetic modulation of NAc-VTA projections. *Related to* Figure 3 *and 4*. **(A)** Left: AAV for Cre-dependent expression of retroChR2 was injected in the NAc (NAcLShell + NAcMShell + NAcCore) of DAT-Cre mice, and multi-unit extracellular recordings coupled with i.v. injection of nicotine were performed. Middle: representative recordings of VTA DA neurons inhibited by 10 Hz optogenetic VTA-NAc activation (ultra-high-power LED: 600-630 nm; 5 min stimulation; *top*) or by i.v. injection of nicotine (*bottom*). Right: responses of VTA DA neurons to 10 Hz stimulation or i.v. nicotine injection. Responses are ranked based on 10 Hz photostimulation and nicotine response, from most activated (red) to most inhibited (pink; n=6 neurons). The horizontal line demarcates neurons showing inhibition (5 out of 6 for light and 4 out of 6 for nicotine) from those showing activation in response to light or nicotine. **(B)** JAWS-expressing AAV was injected in the NAc (NAcLShell + NAcMShell + NAcCore) of WT mice, and multi-unit extracellular recordings coupled with i.v. injection of nicotine were performed. **(C)** Comparison of firing rate variation (Hz) between baseline (Bas) and 5 minutes of continuous optogenetic inhibition (Opto) or saline (Sal) or nicotine (Nic) or optogenetic inhibition coupled with nicotine injection (Opto+Nic) for nicotine-inhibited putative DA neurons (n=6; paired Student’s t-test: Bas vs Opto, t_5_ = -1.12, p = 0.32; Bas vs Sal, t_5_ = 0.8, p = 0.46; Bas vs Nic, t_5_ = 3.55, *p = 0.02; Bas vs Opto+Nic, t_5_ = - 1.3, p = 0.25). **(D)** Example of change in firing rate variation expressed as percentage of baseline after i.v. nicotine injection (Nic) or i.v. nicotine injection coupled with optogenetic inhibition of NAc terminals in the VTA (Opto+Nic). **(E)** Comparison of firing rate variation (Hz) between baseline (Bas) and 5 minutes of continuous optogenetic inhibition (Opto) or saline (Sal) or nicotine (Nic) or optogenetic inhibition coupled with nicotine injection (Opto+Nic) for nicotine-activated putative DA neurons (n = 4; paired Student’s t-test: Bas vs Opto, t_3_ = 0.62, p = 0.58; Bas vs Sal, t_2_ = 4.56, *p = 0.04; Bas vs Nic, t_3_ = -4.55, *p = 0.02; Bas vs Opto+Nic, t_3_ = -3.62, *p = 0.04).

**Figure S6:**
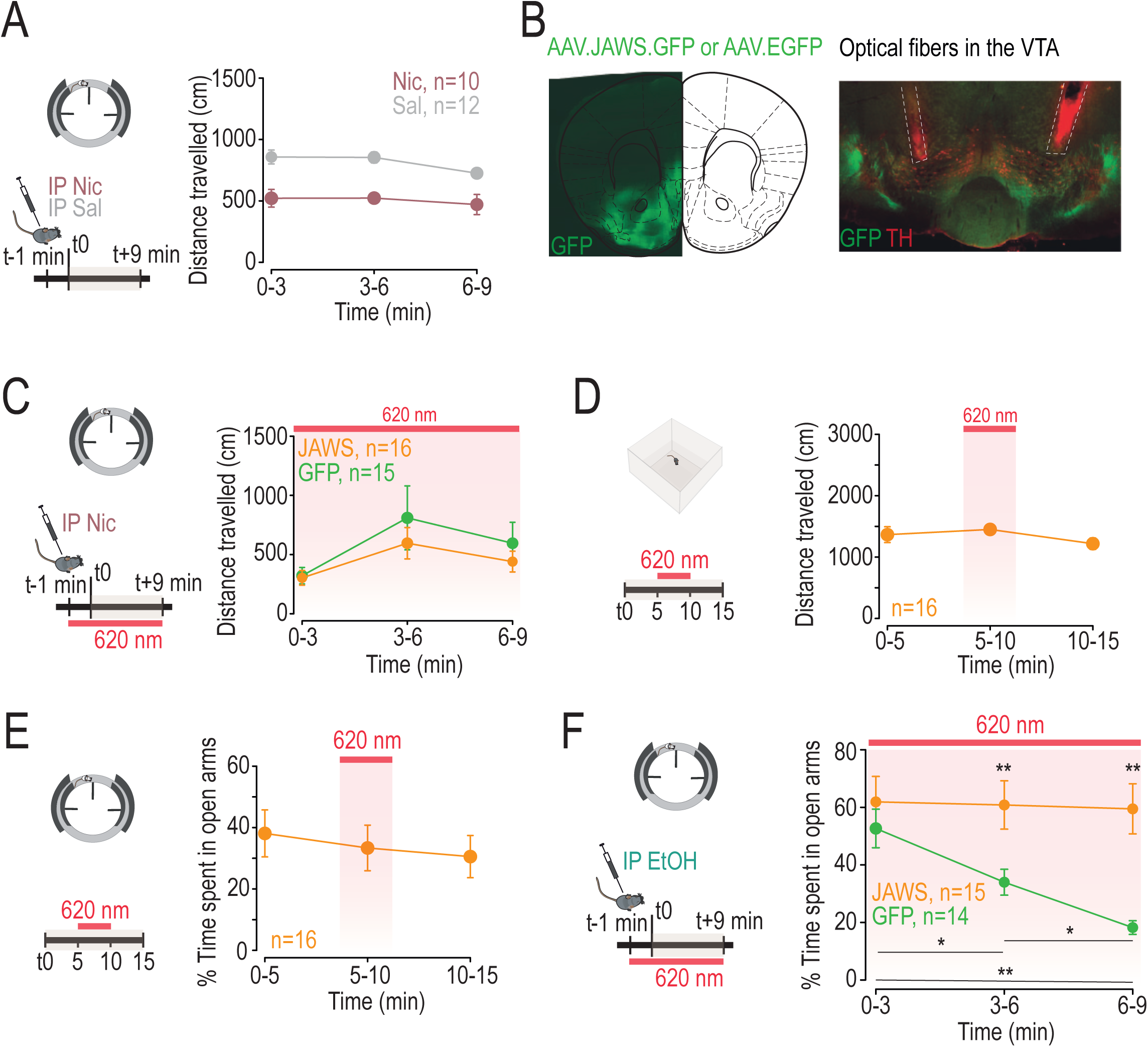
Optogenetic inhibition of NAc terminals in the VTA does not impair locomotion or anxiety-like behavior itself but prevents ethanol-induced anxiety. *Related to* *Figure 4*. **(A)** Locomotor activity in the elevated O maze (EOM) test. Left: intraveneous (i.p.) injection of nicotine (0.5 mg/kg) or saline 1 min before the elevated O maze (EOM) test. Right: distance travelled during the test for saline and nicotine injected mice (n = 10 and 12 respectively; two-way RM ANOVA: group effect, F_1,20_ = 34.31, ***p < 0.001). **(B)** JAWS-expressing AAV was injected in the NAc (NAcLSh, NAcMSh, and NAcCore) and optical fibers were implanted in the VTA. Immunohistochemistry shows viral expression in the NAc (GFP; *left*) and of NAc terminals in the VTA (GFP, TH; *right*). Dashed lines indicate the location of the fibers implanted in the VTA. **(C)** Locomotor activity in the elevated O maze (EOM) test. Left: i.p. nicotine (0.5 mg/kg) was performed 1 min before the EOM test and light stimulation was then continuously maintained throughout the entire test. Right: distance travelled by JAWS- and GFP-expressing mice during the test with continuous inhibition of NAc terminals in the VTA (n = 16 and 15 respectively; two-way RM ANOVA: time effect, F_2,58_ = 5.9, **p = 0.005). **(D)** Locomotor activity was assessed in an open field apparatus. Left: the test lasted 15 minutes and consisted of 5-minute light period (continuous at 620 nm) in between two non-light periods (OFF-ON-OFF). Right: distance travelled of JAWS-expressing mice during the test. No change in locomotor activity was observed upon optogenetic inhibition of NAc terminals in the VTA (n = 16; one-way ANOVA: F_1,46_=0.97, p=0.33). **(E)** Anxiety-like behavior was assessed in the elevated O maze (EOM) test. Left: the test lasted 15 minutes and consisted of 5-minute light period (continuous at 620 nm) in between two non-light periods (OFF-ON-OFF). Right: percentage of time spent in open arms of JAWS-expressing mice during the test. No change in anxiety-like behavior was observed upon optogenetic inhibition of NAc terminals in the VTA (n = 16; one-way ANOVA: F_1,46_=0.54, p=0.46). **(F)** Left: JAWS- or GFP-expressing AAV in the NAc (NAcLShell + NAcMShell + NAcCore) and implantation of optical fibers in the VTA. I.p. ethanol (1 g/kg) was performed 1 min before the EOM test and light stimulation was then continuously maintained throughout the entire test. Right: percentage of the time spent in open arms with continuous inhibition of NAc terminals in the VTA for JAWS-expressing (n = 15) and GFP-control (n = 14) mice (two-way ANOVA: group x time interaction, F_2,54_=4.75, *p=0.01; group effect, F_1,27_=10.02, **p=0.004; time effect, F_2,54_=5.94, **p=0.005; post-hoc Wilcoxon test with Holm corrections for GFP mice: for 3 vs 6 min, V=96, *p=0.04; for 3 vs 9 min, V=102, **p=0.007; for 6 vs 9 min, V=95, *p=0.04; post-hoc Wilcoxon test: JAWS vs GFP at 6 min, W=44, **p=0.008; JAWS vs GFP at 9 min, W=35, **p=0.002; GFP-control group is composed of WT and GFP-expressing mice).

**Figure S7:**
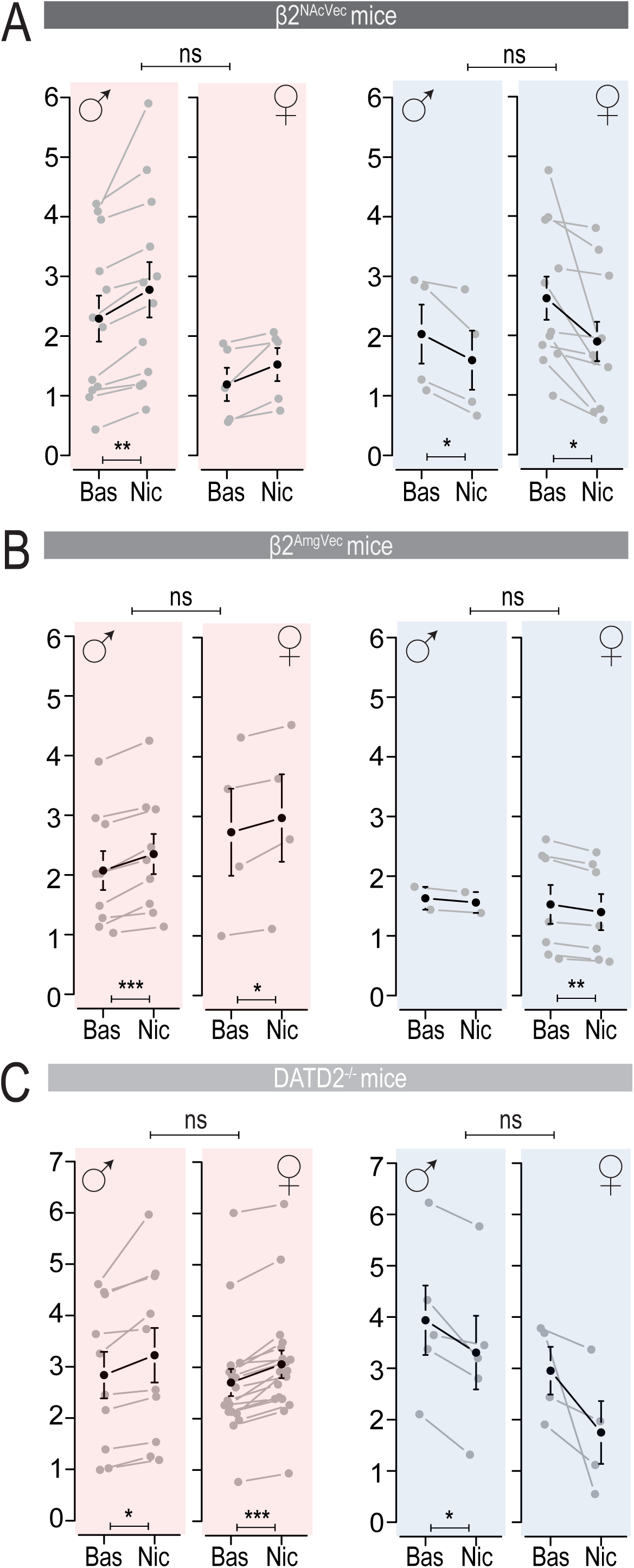
Comparison of nicotine-induced responses *in vivo* between males and females. *Related to methods*. **(A)** Comparison of firing rate variation (Hz) between baseline and nicotine injection and between sexes in β2^NAcVec^ mice. Maximum firing rate for activated neurons (red) or minimum firing rate for inhibited neurons (blue) after i.v. nicotine (Paired t-test for Bas vs Nic: activated neurons in males: t_11_ = -3.7, **p=0.003 or in females t_4_ = -2.56, p = 0.06; inhibited neurons in males: t_3_ = 3.29, *p = 0.046 or in females t_10_ = 2.89, *p = 0.02; Wilcoxon test for comparison between males and females: for activated neurons, W = 39, p = 0.38 or for inhibited neurons, W = 23, p = 0.95). **(B)** Same as (A) in β2^AmgVec^ mice (Paired t-test for Bas vs Nic: activated neurons in males: t_8_ = -5.99, ***p=0.0003 or in females t_3_ = -3.21, *p = 0.049; inhibited neurons in females t_6_ = 4.75, **p = 0.003; Welch two sample t-test or Wilcoxon test for comparison between males and females: for activated neurons, t_5.44_ = 0.43, p = 0.69 or for inhibited neurons, W = 10, p = 0.5). **(C)** Same as (A) in DATD2^-/-^ mice (Paired t-test or Wilcoxon test for Bas vs Nic: activated neurons in males: t_9_ = -3.09, *p = 0.01 or in females V = 0, ***p = 0.0002; inhibited neurons in males t_4_ = 3.6, *p = 0.02; Wilcoxon test for comparison between males and females: for activated neurons, W = 76, p = 0.52 or for inhibited neurons, W = 9, p = 0.9).

## STAR Methods

### LEAD CONTACT

Further information and requests for resources and reagents should be directed to and will be fulfilled by the Lead Contact Fabio Marti

### MATERIALS AVAILABILITY

This study did not generate new unique reagents.

#### Data Availability

All the data that support the findings are available from the corresponding author.

#### Code Availability

All codes used to run the analysis are available from the authors upon request.

## EXPERIMENTAL MODEL

Experiments were performed on wild-type (WT) C57Bl/6Rj (Janvier Labs, France), DATiCRE (DAT-Cre), DAT-D2R^-/-^, ACNB2^-/-^ (β2^-/-^) mice, weighing 25-35 grams. β2^-/-^ mice were provided by the team of Uwe Maskos (Pasteur institute, Paris, France). DATiCRE mice were provided by François Tronche’s team (IBPS Paris, France), and were bred on site and genotyped as described ^35^. DAT-D2R^-/-^ were provided by Emmanuel Valjent’s team (IGF, Montpellier, France). WT and DATiCRE (DAT-Cre) animals were male, whereas DAT-D2R^-/-^ and ACNB2^-/-^ transgenic animals used for in vivo electrophysiology were equally divided between males and females to optimize animal yield. The response of DA neurons to nicotine was similar in both sexes (**Figure S7**).

Mice were kept in an animal facility where temperature (20 ± 2°C) and humidity were automatically monitored and a circadian 12/12-h light-dark cycle was maintained. All experiments were performed on 8-to-20-week-old mice.

All experiments were performed in accordance with the recommendations for animal experiments issued by the European Commission directives 219/1990, 220/1990 and 2010/63, and approved by Sorbonne University and PSL.

## METHODS DETAILS

### Virus

Lentiviruses were prepared in Pasteur institute as previously described ^14,36^, with a titer of either 260 ng/µl for the AChR β2-expressing vector (PDGF.low.mCherry.lox.β2) or 370 ng/µl for GFP-expressing vector (PDGF.low.mCherry.lox.β2). pAAV5-hSyn-hChR2(H134R)-EYFP and pAAV-hsyn-Jaws-KGC-GFP-ER2 were provided by Addgene. ssAAV-retro/2-hSyn1-chI-mCherry_2A_iCre-WPRE-SV40p(A), ssAAV-retro/2-EYFP(rev)-dlox-WPRE-hGHp(A), ssAAV5/2-hSyn1-JAWS-KGC-EGFP-ER2-WPRE-hGHp(A), ssAAV5/2-hSyn1-EGFP-WPRE-hGHp(A) were provided by VVF Zurich.

### Drugs

In all our experiments we used a nicotine hydrogen tartrate salt (Sigma-Aldrich, USA) and liquid ethanol 96% (Emprove, Ph Eur BP, Sigma-Aldrich).

For juxtacellular and tetrode recordings, we performed an intravenous injection (i.v) of nicotine at a dose of 30 µg/kg or ethanol at a dose of 250 mg/kg (and 125, 500, 750 for dose-response experiments) or saline solution (H_2_O with 0.9% NaCl). For patch-clamp recordings, we used 30 µM of nicotine for bath-application and 10, 30 or 100 µM for local puffs. All solutions were prepared in the laboratory.

### Stereotaxic surgeries

For viral or retrobead injections, mice were anesthetized with a gas mixture of oxygen (1 L/min) and 3% isoflurane (IsoFlo) through a TeamSega apparatus. Mice deeply anesthetized were then placed in a stereotaxic frame (David Kopf), maintained under anesthesia throughout the surgery at 3 to 2% isoflurane. The skin was shaved, disinfected and locally anesthetized with 100 µl of lurocaine 10% at the location of the scalp incision. Depending on the experiments, unilateral or bilateral craniotomies were then performed over the VTA, NAc or BLA (see details below). At the end of the surgery, 75 µl of buprenorphine (Buprecare, 0.1 mg/kg) was injected subcutaneously to prepare awakening.

### Retrobead injections

Green or red fluorescent retrograde tracers, retrobeads (RB, Lumafluor), were injected using a cannula (diameter 36G, Phymep, Paris, France). Red RB were used for optogenetic experiments in slice electrophysiology, to not overlap with the YFP signal. Green RB were used for juxtacellular recordings and for patch-clamp recordings without optogenetics. The canula was connected to a 10 µl Hamilton syringe (Mode 1701, Hamilton Robotics, Bonaduz, Switzerland) placed in a pump (QSI, Stoelting Co, Chicago, IL, USA). For juxtacellular recordings, injections were performed in the 3 subregions of the NAc (NAc medial shell NAcMSh: bregma 1.78 mm, lateral 0.45 mm, ventral 4.1 mm; NAc core: bregma 1.55 mm, lateral 1.0 mm, ventral 4.0 mm; NAc lateral shell NAcLSh: bregma 1.45 mm, lateral 1.75 mm, ventral 4.0 mm) or in Amg (BLA: bregma -1.35 mm, lateral 3.07 mm, ventral 4.7 mm; CeA: bregma - 0.78 mm, lateral 2.3 mm, ventral 4.8 mm). For patch-clamp experiments, only NAcLSh (bregma 1.45 mm, lateral 1.75 mm, ventral 4.0 mm) and BLA (bregma -1.35 mm, lateral 3.07 mm, ventral 4.7 mm) were targeted with the same protocol of RB injection. To enable retrograde transport of the RB into the somas of midbrain DA neurons, we waited 2 weeks after injection into the NAc and 3-4 weeks after injection into the Amg, before running electrophysiology experiments.

### Lentiviral reexpression and optogenetic experiments

All viral injections were done using a glass micropipette (10 µl graduated borosilicate glass capillary; Wiretrol I Calibrated Micropipettes, Drummond) prefilled with mineral oil, and fixed into the MO-10 One-axis Oil Hydraulic Micromanipulator (Narishige).

For patch-clamp experiments, activation of NAc terminals was mediated through viral injection of pAAV5-hSyn-hChR2(H134R)-EYFP (2×10^13^, 26973, Addgene).

To perform re-expression of the β2 subunit specifically in the VTA-NAc pathway of β2^-/-^ mice, we first injected ssAAV-retro/2-hSyn1-chI-mCherry_2A_iCre-WPRE-SV40p(A) (8.4×10^12^, v147-retro, VVF Zurich) in the 3 subregions of the NAc (200 nl in each site: NAcMSh: bregma 1.78 mm, lateral 0.45 mm, ventral 4.1 mm; NAcCore: bregma 1.54 mm, lateral 1.0 mm, ventral 4.0 mm; NAcLSh: bregma 1.45 mm, lateral 1.75 mm, ventral 4.0 mm) and we waited 3 weeks for Cre recombinase to expressed. We then performed unilateral injections of 700 nl of PDGF.lox.mCherry.lox.GFP (185 ng/nl, Pasteur institute) or PDGF.lox.mCherry.lox.β2 (260 ng/nl, Pasteur institute) in the VTA (bregma -3.1 mm, lateral 0.5 mm, ventral 4.5 mm).

To perform DA-neuron specific optogenetic activation of the VTA-NAc pathway for patch-clamp or tetrode recordings, we used 8-week-old DAT-Cre mice, in which Cre recombinase expression is restricted to DA neurons without disrupting endogenous dopamine transporter (DAT) expression ^35,37^. We injected ssAAV-retro/2-hEF1a-dlox-hChR2(H134R)_EYFP(rev)-dlox-WPRE-hGHp(A) (5.4×10^12^, v214-retro, VVF Zurich) in the 3 subregions of the NAc (200 nl each site: NAcMSh: bregma 1.78 mm, lateral 0.45 mm, ventral 4.1 mm; NAcCore: bregma 1.55 mm, lateral 1.0 mm, ventral 4.0 mm; NAcLSh: bregma 1.45 mm, lateral 1.75 mm, ventral 4.0 mm) and we waited 3 weeks before recordings.

To perform non-conditional optogenetic inhibition of the NAc inputs onto the VTA for tetrode recordings, we injected pAAV-hsyn-Jaws-KGC-GFP-ER2 (1,3e^13^ 1:10 dilution; 65014; Addgene) or ssAAV5/2-hSyn1-JAWS-KGC-EGFP-ER2-WPRE-hGHp(A) (5,9e^12^; v387-5; VVF Zurich, from #65014 of Addgene) in the 3 subregions of the NAc (200 nl each site: NAcMSh: bregma +1.78 mm, lateral 0.45 mm, ventral 4.1 mm; NAcCore: bregma +1.54 mm, lateral 1.0 mm, ventral 4.0 mm; NAcLSh: bregma +1.45 mm, lateral 1.75 mm, ventral 4.0 mm) in WT mice.

To perform optogenetic behavioral experiments (EOM test, Open field), we did bilateral injections of ssAAV5/2-hSyn1-JAWS-KGC-EGFP-ER2-WPRE-hGHp(A) (5,9e^12^; v387-5; VVF Zurich, from #65014 of Addgene) or ssAAV5/2-hSyn1-EGFP-WPRE-hGHp(A) (5,7e^12^; v81-5; VVF Zurich) in the NAc (200 nl each site: NAcMSh: bregma +1.78 mm, lateral 0.45 mm, ventral 4.1 mm; NAcCore: bregma +1.54 mm, lateral 1.0 mm, ventral 4.0 mm; NAcLSh: bregma +1.45 mm, lateral 1.75 mm, ventral 4.0 mm) in WT mice. Optical fibers (200 mm core, NA = 0.39, ThorLabs) coupled to a zirconia ferule (1.25 mm) were implanted bilaterally in the VTA (10° angle; bregma -3.07 mm, lateral 1.43 mm, ventral -3.9 mm) and fixed to the skull with dental cement (SuperBond, Sun medical).

### *In vivo* electrophysiology on anesthetized animals

Induction of anesthesia was done with gas mixture of oxygen (1 L/min) and 3% isoflurane (IsoFlo) through a TeamSega apparatus. Mice deeply anesthetized were then placed in a stereotaxic frame (David Kopf), maintained under anesthesia throughout the surgery at 3 to 2.5% isoflurane. The scalp was opened and a cranial window was drilled in the skull above the location of the VTA (coordinates: 3.1 ± 3 mm posterior to bregma, 0.4 to 0.5 mm lateral to the midline, 3.9 to 5 mm ventral from the brain). During recordings, mice were maintained deeply anesthetized at 2 % isoflurane, with monitoring and adjustment of the anesthesia throughout the experiment. Intravenous (i.v.) administration of saline, nicotine (30µg/kg) or ethanol (250 mg/kg) was carried out through a catheter (30G needle connected to polyethylene tubing PE10) connected to a Hamilton syringe, into the saphenous vein of the animal. For multiple doses of ethanol, mice received two to four injections of 125, 250, 500 and/or 750 mg/kg (pseudo-randomly administrated).

### Juxtacellular recordings

To perform single unit extracellular recordings, recording electrodes were pulled from borosilicate glass capillaries (Harvard Apparatus, with outer and inner diameters of 1.50 and 1.17 mm, respectively) with a Narishige electrode puller. The tips were broken under microscope control and filled with 0.5% sodium acetate containing 1.5% of neurobiotin tracer (VECTOR laboratories). Electrodes had tip diameters of 1-2 µm and impedances of 6–9 MΩ. A reference electrode was placed in the subcutaneous tissue. The recording electrodes were lowered vertically through the hole with a micro drive. Electrical signals were amplified by a high-impedance amplifier (Axon Instruments) and monitored through an audio monitor (A.M. Systems Inc.). The unit activity was digitized at 25 kHz and recorded using Spike2 software (Cambridge Electronic Design) for later analysis. Individual electrode tracks were separated from one another by at least 0.1 mm in the medio-lateral axis. The electrophysiological characteristics of dopamine neurons were analyzed in the active cells encountered when passing the microelectrode in a stereotaxically defined block of brain tissue corresponding to the coordinates of the VTA (coordinates: between 3 to 3.4 mm posterior to bregma, 0.4 to 0.6 mm lateral to midline, and 3.9 to 5 mm below brain surface). Extracellular identification of dopamine neurons was based on their location as well as on the set of unique electrophysiological properties that distinguish dopamine from non-dopamine neurons *in vivo*: (i) a typical triphasic action potential with a marked negative deflection; (ii) a long duration (>2.0 ms); (iii) an action potential width from start to negative trough >1.1 ms; (iv) a slow firing rate (<10 Hz and >1 Hz). After recording, nicotine-responsive cells were labeled by electroporation of their membrane: successive currents squares were applied until the membrane breakage, to fill cell soma with neurobiotin contained into the glass pipet ^38^. To be able to establish correspondence between neurons responses and their localization in the VTA, we labeled one type of response per mouse: solely activated neurons or solely inhibited neurons, with a limited number of cells per brain (1 to 4 neurons maximum, 2 by hemisphere), always with the same concern of localization of neurons in the VTA. All data were analyzed with R.

### Multi-unit extracellular recordings

4-5 weeks after viral infection, we used a Mini-Matrix (Thomas Recording) allowing us to lower within the VTA up to 3 tetrodes (Tip shape A, Thomas Recording, Z = 1–2 MW) and a tip-shaped quartz optical fiber (100 mm core, NA = 0.22, Thomas Recording) for photostimulation. The fiber was coupled to a dual LED (450-465 nm for ChR2, 600-630 nm for Jaws, Prizmatix) with an output intensity of 200–500 mW for both wavelengths. These four elements could be moved independently with micrometer precision. Tetrodes were lowered in the medial part of the VTA while the optical fiber was lowered in the lateral part of the VTA 100 μm above the tetrodes (around 300 μm between the fiber and the tetrodes).

Spontaneously active putative DA neurons were identified on the basis of the same electrophysiological criteria used for juxtacellular recordings. Baseline activity was recorded for 5 minutes, prior to i.v. injection of nicotine, allowing us to target drug-inhibited neurons and a second i.v. injection of drug with optical-stimulation was then performed.

Electrophysiological signals were acquired with a 20 channels preamplifier included in the Mini Matrix (Thomas Recording) connected to an amplifier (Digital Lynx SX 32 channels, Neuralynx) digitized and recorded using Cheetah software (Neuralynx). Spikes were detected (CSC Spike Extractor software, Neuralynx) and sorted using a classical principal component analysis associated with a cluster cutting method (SpikeSort3D Software, Neuralynx). All the data were analyzed with R.

### *Ex vivo* electrophysiology: patch clamp recordings

Mice were deeply anesthetized by an intraperitoneal injection of a mix of ketamine (150 mg/kg Virbac 1000) and xylazine (60 mg/kg, Rompun 2%, Elanco). Coronal midbrain sections (250 µm) were sliced with a Compresstome (VF-200, Precisionary Instruments) after intracardial perfusion of cold (4°C) sucrose-based artificial cerebrospinal fluid containing (in mM): 125 NaCl, 2.5 KCl, 1.25 NaH_2_PO_4_, 26 NaHCO_3,_ 5,9 MgCl_2_, 25 sucrose, 2.5 glucose, 1 kynurenate (pH 7.2, 325 mOsm). After 8 minutes at 37°C for recovery, slices were transferred into oxygenated artificial cerebrospinal fluid (ACSF) containing (in mM): 125 NaCl, 2.5 KCl, 1.25 NaH_2_PO_4_, 26 NaHCO_3,_ 2 CaCl2, 1 MgCl2, 15 sucrose, 10 glucose (pH 7.2, 325 mOsm) at room temperature for the rest of the day. Slices were individually transferred to a recording chamber continuously perfused at 2 mL/minute with oxygenated ACSF. Patch pipettes (4-6 MΩ) were pulled from thin wall borosilicate glass (G150TF-3, Warner Instruments) with a micropipette puller (P-87, Sutter Instruments Co.). Neurons were visualized using an upright microscope coupled with a Dodt gradient contrast imaging, and illuminated with a white light source (Scientifica). Whole-cell recordings were performed with a patch-clamp amplifier (Axoclamp 200B, Molecular Devices) connected to a Digidata (1550 LowNoise acquisition system, Molecular Devices). Signals were low-pass filtered (Bessel, 2 kHz) and collected at 10 kHz using the data acquisition software pClamp 10.5 (Molecular Devices). VTA location was identified under microscope. Identification of dopaminergic neurons was performed by location and by their electrophysiological properties (width and shape of action potential (AP) and after hyperpolarization (AHP).

To perform recordings of spontaneous inhibitory post-synaptic currents (sIPSCs), we used a Cesium-based internal solution of (in mM): 130 CsCl, 1 EGTA, 10 HEPES, 2 MgATP, 0.2 NaGTP, 0.1% neurobiotin pH 7.35 (270-285 mOsm). Local perfusion was used to apply nicotine locally in the bath above the recorded cells. Recordings of light-evoked GABA current from activation of NAc terminals were conducted in the presence of 20 µM DNQX (6,7-Dinitroquinoxaline-2,3-dione, HelloBio) and 50 µM D-AP5 (D-(-)-2-Amino-5-phosphonopentanoic acid, HelloBio) to block respectively AMPA and NMDA receptors, 500 mM of TTX (tetrodotoxin citrate, HelloBio) to block voltage-gated sodium channels and 1mM of 4-AP (4-aminopyridine, HelloBio) to block voltage-gated potassium channels ^39^. Light-evoked responses were obtained every 10s with one pulse (3ms) of 460 nm wavelength (5 sweeps for each neuron). For recordings of miniature inhibitory post-synaptic currents (mIPSCs), Cs-based internal solution was used with 20 µM DNQX, 50 µM D-AP5 and 500 nM TTX.

To perform characterization of DA subpopulations and puffs recordings (200 ms, 2 psi), potassium gluconate-based intracellular solution was used containing (in mM): 135 K-gluconate, 10 HEPES, 0.1 EGTA, 5 KCl, 2 MgCl_2_, 2 ATP-Mg, 0.2 GTP-Na, and biocytin 2 mg/mL (pH adjusted to 7.2). The same internal solution was used to record optical stimulation of retroChR2 virus in NAc-projecting DA neurons. To characterize retroChR2 expression, 10 and 20 Hz (3s pulse; train rate of 10 or 20 Hz; 5 sweeps of 10s) photostimulation were used to drive neuronal firing in current-clamp mode. LED (Prizmatix) intensity was set to 30%, as no higher amplitude currents were observed above this threshold.

### Behavioral task

#### Elevated O maze test

The raw data for behavioral experiments were acquired as video files. The elevated O-maze (EOM) apparatus consists of two open (stressful) and two enclosed (protecting) elevated arms that together form a zero or circle (diameter of 50 cm, height of 58 cm, 10 cm-wide circular platform). Time spent in exploring enclosed versus open arms indicates the anxiety level of the animal. The first EOM experiment assessed the effect of an i.p. injection of Nic (0.5 mg/kg) on WT mice. The test lasts 10 minutes: mice are injected 1 minute before the test, and then put in the EOM for 9 minutes. In the second EOM experiment, i.p. injection of Nic (0.5 mg/kg) or Ethanol (1 g/kg) was combined with continuous optogenetic inhibition of NAc terminals targeting the VTA. Finally, we did an optogenetic EOM experiment to control for basal anxiety-like behavior under light-stimulation, during 15 min, alternating 5 minute-periods of stimulation and non-stimulation (OFF-ON-OFF). Time spent in open or closed arms was extracted frame-by frame using the open-source video analysis pipeline ezTrack ^40^. Mice were habituated to the stress of handling and injection for a minimum of one week before testing.

#### Open field test

The open field (OF) is a square enclosure of 50 cm x 50 cm where animals can move freely. Distance travelled by animals was quantified over time. Regarding the optogenetic experiment conducted in the OF, animals were placed in the maze for 15 minutes, while alternating between OFF, ON and OFF optical stimulation periods of 5 minutes each.

### Fluorescence immunohistochemistry

Recorded neurons in patch-clamp experiments were filled with biocytin in order to validate the presence of TH enzyme by immunohistochemistry, indicator of dopaminergic neurons. After recordings, slices were fixed in 4% PFA (paraformaldehyde) during a night. Recorded neurons in juxtacellular expriments were filled with neurobiotin and after euthanasia of the animals, brains were rapidly removed and fixed in 4% PFA. After a period of at least three days of fixation at 4°C, serial 60-μm sections were cut from the midbrain with a vibratome.

Immunostaining experiments were performed as follows: free-floating VTA brain sections were incubated for 3 hours (juxtacellular experiments) or 6 hours (patch-clamp experiments) at 4°C in a blocking solution of phosphate-buffered saline (PBS) containing 3% bovine serum albumin (BSA, Sigma; A4503) and 0.2% Triton X-100 (vol/vol), and then incubated overnight (juxtacellular experiments) or during 72 hours (patch-clamp experiments) at 4°C with a mouse anti-tyrosine hydroxylase antibody (anti-TH, Sigma, T1299), at 1:500 or 1:200 dilution, in PBS containing 1.5% BSA and 0.2% Triton X-100. Sections were then rinsed with PBS, and incubated for 3 hours (juxtacellular experiments) or 6 hours (patch-clamp experiments) at room temperature with Cy3-conjugated anti-mouse and AMCA-conjugated streptavidin (Jackson ImmunoResearch) both at 1:500 or 1:200 dilution in a solution of 1.5% BSA in PBS. After three rinses in PBS, slices were wet-mounted using Prolong Gold Antifade Reagent (Invitrogen, P36930). In the case of optogenetic or re-expression experiments, identification of transfected neurons by immunofluorescence was performed as described above, with addition of chicken anti-GFP primary antibody (1:500, ab13970, Abcam) in the 3% BSA solution. A goat-anti-chicken AlexaFluor 488 (1:500, Life Technologies) was then used as secondary antibody.

Microscopy was carried out with a fluorescent microscope, and images captured using a camera and analyzed with ImageJ. Immunoreactivity for both TH and biocytin or neurobiotin allowed us to confirm the neurochemical phenotype of DA neurons in the VTA or the transfection success.

### Quantification and analysis of *in vivo* electrophysiological recordings

Subpopulations of DA neurons were automatically classified using variation of firing frequency induced by nicotine or ethanol i.v injection. First, we calculated the maximal variation from the baseline per neuron, within the first 3 minutes following injection. We then used a bootstrapping method to exclude non-responding neurons.

For each neuron, the maximal and the minimal value of firing frequency was measured within the response period (3 minutes) that followed nicotine or saline injection, and the biggest of the two were used to determine the maximum of variation induce by the given injection. The effect of nicotine or alcohol was assessed by comparing the maximum of firing frequency variation induced by the drug and by saline injection. If the distribution of all maximum variation induced by nicotine or ethanol were different from those induced by saline, each neuron was then individually classified as activated or inhibited by bootstrapping. Baseline spike intervals were randomly shuffled 1000 times. Firing frequency was estimated on 60s-time windows, with 15 s time steps. For each neuron, we determined the percentile from the shuffled data corresponding to the drug-evoked response (max or min frequency after nicotine injection). Neurons were individually considered as responsive to nicotine or ethanol injection if this percentile is ≥ 0.95 or ≤ 0.05. Responsive neurons displaying an increase in firing frequency (Δf > 0) were defined as ‘‘Nic+’’ or “EtOH+” while neurons displaying a decrease in firing frequency (Δf < 0) were defined as ‘‘Nic-’’ or “EtOH-“. For the dose-response curve, neurons were classified as EtOH+ or EtOH-based on their response for the first doses inducing a significant firing variation identified by bootstrapping.

The mean responses to nicotine and ethanol of activated and inhibited neurons are thus presented as a percentage of variation from baseline (mean ± SEM). Then, for activated (or inhibited) neurons, we compare the maximum (or minimum) value of firing frequency before and after injection for nicotine, ethanol, and saline. Finally, to ensure that the drug-induced responses were different from the activation or inhibition induced by saline injection, we compared the firing variation induced by nicotine or ethanol with that induced by saline

To map the locations of EtOH+ and EtOH-neurons on an atlas, we included all neurons identified as ethanol-responsive by bootstrapping, regardless of whether they received nicotine or saline injections.

To examine the correlation between ethanol- and nicotine-induced responses, we included all neurons identified as ethanol-responsive by bootstrapping that received a nicotine injection, regardless of whether they received saline injection or not.

### Statistical analysis

All statistical analyses were done using the R software with home-made routines. Results were plotted as mean ± SEM. The total number (n) of observations in each group and the statistical tests used for comparisons between groups or within groups are indicated directly on the figures or in the results sections. Comparisons between means were performed with parametric tests such as Student’s t-test, or two-way ANOVA for comparing two groups when parameters followed a normal distribution (Shapiro-Wilk normality test with p > 0.05), or Wilcoxon non-parametric test if the distribution was skewed. Holm’s sequential Bonferroni post hoc analysis was applied, when necessary. Statistical significance was set at p < 0.05 (*), p < 0.01 (**), or p < 0.001 (***), or p > 0.05 was considered not to be statistically significant. For the surrogate analyses, nicotine and ethanol responses from the experimental data groups were pooled to reconstruct a new surrogate data set. The percentage of correlation of drug responses from the surrogate was then calculated (number of pairs of responses showing the same polarity for the two drugs). We plotted the density of the percentage correlation from 10,000 surrogates and counted the number of times this percentage reached the level of the percentage of correlation observed with experimental data.

### Statistics and Reproducibility

All experiments were replicated with success.

